# Genome-wide CRISPR Screens in Primary Human T Cells Reveal Key Regulators of Immune Function

**DOI:** 10.1101/384776

**Authors:** Eric Shifrut, Julia Carnevale, Victoria Tobin, Theodore L. Roth, Jonathan M. Woo, Christina Bui, P. Jonathan Li, Morgan Diolaiti, Alan Ashworth, Alexander Marson

## Abstract

Human T cells are central effectors of immunity and cancer immunotherapy. CRISPR-based functional studies in T cells could prioritize novel targets for drug development and improve the design of genetically reprogrammed cell-based therapies. However, large-scale CRISPR screens have been challenging in primary human cells. We developed a new method, <u>s</u>gRNA <u>l</u>entiviral <u>i</u>nfection with <u>C</u>as9 protein <u>e</u>lectroporation (SLICE), to identify regulators of stimulation responses in primary human T cells. Genome-wide loss-of-function screens identified essential T cell receptor signaling components and genes that negatively tune proliferation following stimulation. Targeted ablation of individual candidate genes validated hits and identified perturbations that enhanced cancer cell killing. SLICE coupled with single-cell RNA-Seq revealed signature stimulation-response gene programs altered by key genetic perturbations. SLICE genome-wide screening was also adaptable to identify mediators of immunosuppression, revealing genes controlling response to adenosine signaling. The SLICE platform enables unbiased discovery and characterization of functional gene targets in primary cells.

## INTRODUCTION

Cytotoxic T cells play a central role in immune-mediated control of cancer, infectious diseases, and autoimmunity. Immunotherapies such as checkpoint inhibitors and engineered cell-based therapies are revolutionizing cancer treatments, achieving durable responses in a subset of patients with otherwise refractory malignant disease (June et al., 2018; Reck et al., 2016; Wolchok et al., 2017). However, despite dramatic results in some patients, the majority of patients do not respond to available immunotherapies (Sharma et al., 2017). Much work remains to be done to extend immunotherapy to common cancers that remain refractory to current treatments.

Next-generation adoptive cell therapies are under development utilizing CRISPR-Cas9 genome engineering. Cas9 ribonucleoproteins can be delivered to primary human T cells to efficiently knockout checkpoint genes (Ren et al., 2017a; Rupp et al., 2017; Schumann et al., 2015) or even re-write endogenous genome sequences (Roth et al., 2018). While deletion of the canonical checkpoint gene encoding PD-1 may enhance T cell responses to some cancers (Ren et al., 2017b; Rupp et al., 2017), an expanded set of targets would offer additional therapeutic opportunities. Advances in immunotherapy depend on improved understanding of the genetic programs that determine how T cells respond when they encounter their target antigens. Promising gene targets could enhance T cell proliferation and productive effector responses upon stimulation. In addition, immunosuppressive cells and soluble molecules such as cytokines and metabolites can accumulate within tumors and hamper productive anti-tumor T cell responses. Gene targets that influence a T cell’s ability to overcome immunosuppressive tumor microenvironments could extend the reach of adoptive cell therapies to solid tumors.

Decades of work in animal models and cell lines have identified regulators of T cell suppression and activation, but systematic strategies to comprehensively analyze the function of genes that regulate human T cell responses are still lacking. Gene knock-down with curated RNA interference libraries have pointed to targets that enhance *in vivo* antigen-responsive T cell proliferation in mouse models (Zhou et al., 2014). More recently, CRISPR-Cas9 has ushered in a new era of functional genetics studies (Doench, 2018). Large libraries of single guide RNAs (sgRNAs) are readily designed to target genomic sequences. Transduction of cells with lentivirus encoding these sgRNAs generates pools of cells with diverse genomic modifications that can be tracked by sgRNA sequences in integrated provirus cassettes. This approach has been used in cell lines engineered to express stable Cas9 and in Cas9 transgenic mouse models (Parnas et al., 2015; Shang et al., 2018). Pooled CRISPR screens are already revealing gene targets in human cancer cells that modulate responses to T cell-based therapies (Manguso et al., 2017; Pan et al., 2018; Patel et al., 2017). However, CRISPR screening in primary human T cells – which can only be cultured *ex vivo* for limited time spans – has been hampered by low lentiviral transduction rates with Cas9-encoding vectors (Seki and Rutz, 2018). Genome-scale CRISPR screens in human T cells would enable comprehensive studies that may reveal targets that could be rapidly translated into new immunotherapies with small molecules, biologics, and gene-engineered adoptive cell therapies.

Here we developed a screening platform that combines pooled lentiviral sgRNA delivery with Cas9 protein electroporation to enable loss-of-function pooled screening at genome-wide scale in primary human T cells. We applied this technology to identify gene modifications that promote T cell proliferation in response to stimulation. A subset of hits from these screens enhanced *in vitro* anti-cancer activity of human T cells, suggesting that they may be promising preclinical candidates for next-generation cell therapies. We further coupled pooled CRISPR delivery with single-cell transcriptome analysis of human T cells to characterize the cellular programs controlled by several of the genes found to regulate T cell responses in our genome-wide screens. Finally, we adapted the genome-wide screening context to model suppression by a well-described immunosuppressive metabolite, adenosine, to identify known as well as novel targets that enable escape from adenosine receptor-mediated immunosuppression. Taken together, these studies provide a rich resource of gene pathways that can be targeted to tune human T cell responses and a broadly applicable platform to probe primary human T cell biology at genome-scale.

## RESULTS

### A Hybrid Approach to Introduce Traceable Genetic Perturbations in Primary Human T Cells

We set out to establish a high-throughput CRISPR screening platform that works directly in *ex vivo* human hematopoietic cells. Current pooled CRISPR screening methods rely on establishing cell lines with stably integrated Cas9 expression cassettes. Our attempts to stably express *Streptococcus pyogenes* Cas9 by lentivirus in primary T cells resulted in extremely poor transduction efficiencies. This low efficiency was prohibitive of large-scale pooled screens with primary cells, which are not immortalized and can only be expanded in culture for a limited amount of time. We previously showed efficient gene editing of primary human T cells by electroporation of Cas9 protein pre-loaded *in vitro* with sgRNAs (Hultquist et al., 2016; Schumann et al., 2015). We conceived of a hybrid system to introduce traceable sgRNA cassettes by lentivirus followed by electroporation with Cas9 protein (**Figure 1A**). To test this strategy, we targeted the gene encoding a candidate cell surface protein, the alpha chain of the CD8 receptor (*CD8A*), as it is highly and uniformly expressed in human CD8^+^ T cells. We optimized multiple steps in lentiviral transduction, Cas9 electroporation, and T cell stimulation to ensure efficient delivery of each component while maintaining cell viability and proliferative potential (**Figure S1A-S1D**). Briefly, CD8^+^ T cells were isolated from peripheral blood of healthy donors, stimulated, and then transduced with lentivirus encoding an sgRNA cassette and an mCherry fluorescence protein reporter gene. Following transduction, T cells were transfected with recombinant Cas9 protein by electroporation. At day 4 post-electroporation, the transduced cells (mCherry+) were largely (>80%) CD8 negative (**Figure 1B** and **Figure S1E**), indicative of successful targeting by the Cas9-sgRNA combination. Loss of CD8 protein was specifically programmed by the targeting sgRNA, as cells transduced with a non-targeting control sgRNA retained high levels of CD8 expression. By targeting *PTPRC* (CD45) with the same delivery strategy, we confirmed successful knockout at a second target and demonstrated efficacy of the system in both CD8^+^ and CD4^+^ T cells (**Figure S1F**). We conclude that sgRNA lentiviral infection with <u>C</u>as9 protein <u>e</u>lectroporation (SLICE) results in effective and specific disruption of target genes.

**Figure 1.**
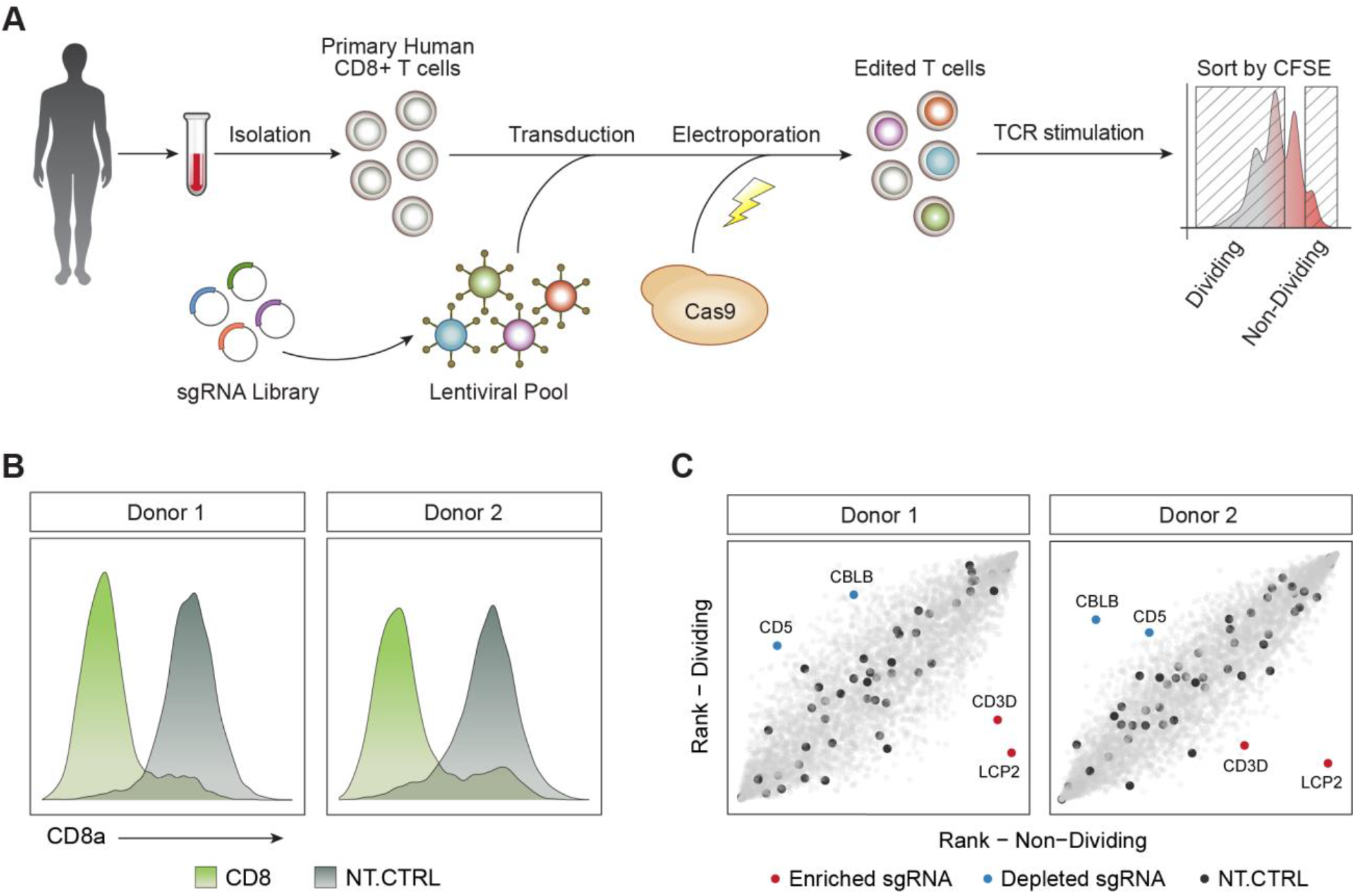
Framework for Unbiased Discovery of Regulators of Human T Cell Proliferation Using Pooled CRISPR Screens. (A) Diagram of a hybrid system of sgRNA lentiviral infection and Cas9 electroporation (SLICE), enabling pooled CRISPR screens in primary human T cells. (B) Editing of the *CD8A* gene with SLICE led to efficient protein knockdown in two independent donors. (C) Targeted screen (4,918 guides) shows that sgRNAs targeting *CBLB* and *CD5* were enriched in proliferating T cells (blue), while sgRNAs targeting *LCP2* and *CD3D* were depleted (red). Non-targeting sgRNAs were evenly distributed across the cell populations (black). See Also Figure S1 and Table S2.

We next tested whether SLICE could be expanded to allow large-scale loss-of-function screens in primary cells with pools of lentivirus-encoded sgRNAs. We performed a screen to identify gene targets that regulate T cell proliferation in response to T cell receptor (TCR) stimulation. For pilot studies we generated a custom library of sgRNA plasmids targeting all annotated cell surface proteins and several canonical members of the TCR signaling pathway (4,918 guides targeting 1211 genes total and 48 non-targeting guides, **Table S1**). CD8^+^ T cells isolated from two healthy human donors were transduced with lentivirus encoding this sgRNA library, electroporated with Cas9, and then maintained in culture (**Experimental Procedures**). At day 10 postelectroporation, cells were labeled with CFSE to track cell divisions and then TCR stimulated. After four days of stimulation, CFSE levels revealed that the cells had undergone multiple divisions. Cells were sorted by FACS into two populations: (1) non-proliferating cells (CFSE high) and (2) highly-proliferating cells (CFSE low) (**Figure 1A**, **Figure S1G**, and **Experimental Procedures**). We quantified sgRNA abundance from each population by deep sequencing of the amplified sgRNA cassettes. Consistent with well-maintained coverage of sgRNAs across experimental steps, we were able to detect all library guides in the infected CD8^+^ T cells, with the distribution of sgRNA abundance being relatively uniform for each donor and across biological replicates (**Figure S1H**). To identify sgRNAs that regulated T cell proliferation, we calculated the abundance-based rank difference between the highly dividing cells and non-dividing cells. sgRNAs with strong enrichment in dividing or non-dividing cells pointed to key biologic pathways. We found that sgRNAs targeting essential components of TCR signaling such as *CD3D* and *LCP2*, inhibited cell proliferation (de Saint Basile et al., 2004; Shen et al., 2009). We also found that proliferation could be enhanced in human T cells by targeting *CD5* or *CBLB*, which have reported roles in negative regulation of T cell stimulation-responses (Azzam et al., 2001; Naramura et al., 2002; Voisinne et al., 2016). sgRNAs targeting these genes were in the top 1% by rank difference in both biological replicates (**Figure 1C**). Furthermore, multiple sgRNAs targeting these genes had concordant effects, increasing our confidence that the phenotype was not due to off-target effects (**Figure S1I**, **Table S2**). Importantly, sorting dividing and non-dividing primary cells based on CFSE provided much stronger enrichment of sgRNA sequences than simple growth-based screens with otherwise identical experimental timelines (**Figure S1J**). Growth-based screens have been largely successful using immortalized cell lines that can be cultured for prolonged durations, however this did not translate to screens in primary human T cells (Shalem et al., 2014; Wang et al., 2014). Taken together, these data demonstrate that SLICE pooled CRISPR screens can be used to discover positive and negative regulators of proliferation in primary human T cells.

### A Genome-wide Pooled CRISPR Screen Uncovers Regulators of the TCR Response

Pooled screens enable high-throughput screening capacity. To take full advantage of this platform, we scaled-up from the targeted pilot screen to genome-wide (GW) scale (Doench et al., 2016), transducing a library of 77,441 sgRNAs (19,114 genes) into T cells from two healthy donors. After confirming successful transduction of these primary human T cells (**Figure S2A-B**), the cells were re-stimulated and then FACS sorted into non-proliferating and highly-proliferating populations based on CFSE levels (**Figure S2C** and **Experimental Procedures**). MAGeCK software (Li et al., 2014) was used to systematically identify genes that are positively or negatively selected in the proliferating population of T cells (**Table S3**). Top positive and negative regulators from the pilot screen were confirmed in both biological replicates of the GW screen along with numerous other hits (**Figure 2A, B**). To hone the list of top candidates, we performed an independent secondary screen in cells from two additional human blood donors. The results were well correlated between the primary and secondary screens (**Figure 2C**). Integrated analysis of the two independent screens performed on a total of four human blood donors provided improved power for target discovery, particularly for negative regulators of T cell proliferation (**Table S4** and **Figure S2D**). To confirm that the hits were in fact dependent on TCR stimulation, we performed GW screens with increasing levels of TCR stimulation. While similar gene targets appeared as positive and negative regulators across the conditions, the magnitude of the effects were blunted at higher levels of TCR stimulation, suggesting that stronger TCR stimulation can override the effects of these genetic perturbations (**Figure 2D** and **Figure S2E**). The observed dose-response confirmed that the majority of the screen hits are dependent on the TCR stimulus and serve to tune resulting proliferative responses. Taken together these screens identified dozens of genetic perturbations that positively and negatively modulate T cell proliferation.

**Figure 2.**
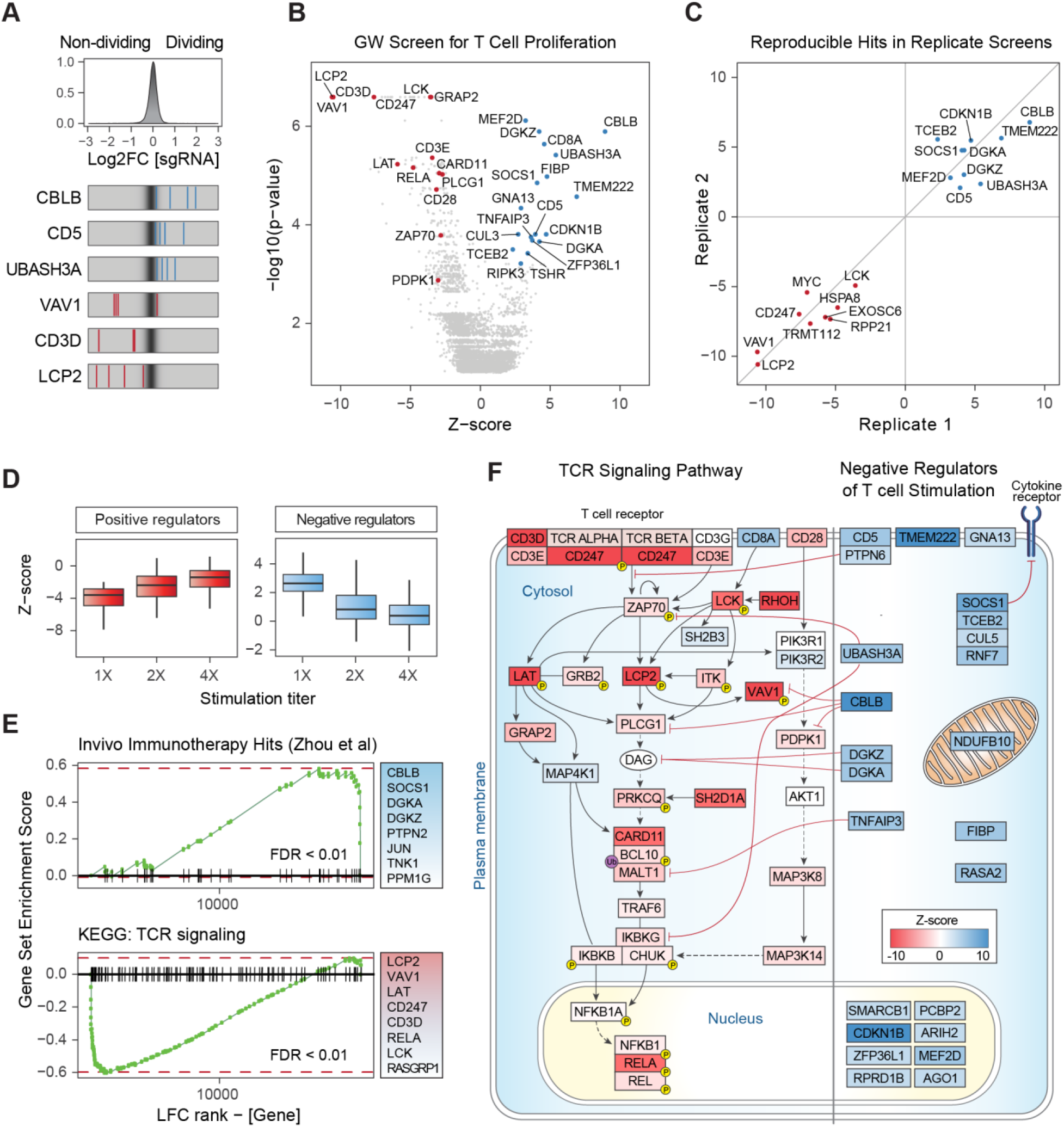
Genome-wide Screen in Primary Human T Cells Identifies Key Mediators of TCR Signaling Dependent T Cell Proliferation. (A) Top panel: distribution of log2 fold-change (LFC) values of dividing over non-dividing cells for >75,000 guides in the genome-wide (GW) library. Bottom panel: LFC for all four sgRNAs targeting three genes enriched in dividing cells (blue lines) and three depleted genes (red lines), overlaid on grey gradient depicting the overall distribution. Values are averaged over two donors. (B) Volcano plot for hits from the primary GW screen. X-axis shows Z-score (ZS) for gene-level LFC (median of LFC for all sgRNAs per gene, scaled). Y-axis shows the p-value as calculated by MAGeCK. Highlighted in red are negative hits (depleted in dividing cells, FDR < 0.2 and |ZS| > 2), which are annotated for the TCR signaling pathway by Gene Ontology (GO). Blue dots show all positive hits (Rank < 20 and |ZS| > 2). All values are calculated for two donors as biological replicates. (C) Gene hits from the secondary GW screen in cells from two independent blood donors are positively correlated with the hits from the primary screen. Shown are Z-scores for overlapping hits for the top 25 ranking targets from the independent screens, in both positive and negative directions. (D) Boxplot for Z-scores (scaled LFC) of the top 100 hits in each direction, for three GW screens with increasing TCR stimulation levels (1X = data in (B)). For both panels, LFC values trended towards 0, indicating selection pressure was reduced as the TCR signal increased. Horizontal line is the median, vertical line is the data range. (E) Gene-set enrichment analysis shows significantly skewed LFC ranking of screen hits in two curated gene lists: (top panel) previously discovered hits by an shRNA screen in a mouse model of melanoma (Zhou et al., 2014) and (bottom panel) TCR signaling pathway by KEGG. The top eight gene members on the leading edge of each set enrichment are shown in the text-box on the right. Vertical lines on the x-axis are members of the gene set, ordered by their LFC rank in the GW screen. FDR = False discovery rate, permutation test. (F) Modulators of TCR signaling and T cell activation detected in the GW screens. Depicted on the left are positive regulators of the TCR pathway found in our GW screens (FDR < 0.25). The curated TCR pathway is based on NetPath_11 (Kandasamy et al., 2010) and literature review. Depicted on the right are negative regulator genes (both known and unknown) found in our GW screens (FDR < 0.25), and represent candidate targets to boost T cell proliferation. Cellular localization and interaction edges are based on literature review. Gene nodes are shaded by their Z-score in the GW screen (red for positive and blue for negative Z-score values). See Also Figure S2 and Tables S3-4.

Genes identified in the integrated screen analysis were enriched with annotated pathways associated with TCR stimulation. Gene set enrichment analysis (GSEA) revealed significant over-representation of gene targets depleted from proliferating cells in the TCR signaling pathway (FDR < 0.01, **Figure 2E and Figure S2F**). We also found a strong enrichment of genes abundant in dividing cells for hits from a published shRNA screen (Zhou et al., 2014) designed to discover gene targets that boost T cell proliferation in tumor tissue *in vivo* (FDR < 0.01). This result is striking, as the studies were done in a different organism with a different gene perturbation platform, yet there was significant enrichment for high ranking positive hits in our screen with the hit list discovered in this *in vivo* animal model. These global analyses confirmed that our functional screens could identify critical gene targets, now achievable at genome-wide scale directly in primary human cells.

Targets depleted from proliferating cells in this GW screen encode key protein complexes critical for TCR signaling (**Figure 2F**). For example, gene targets that impaired TCR dependent proliferation included the delta and zeta chains of the TCR complex itself (negative rank 18 and 6, respectively), LCK (negative rank 20) which directly phosphorylates and activates the TCR ITAMs and the central signaling mediator ZAP70 (negative rank 299) (Dave et al., 1998; Tsuchihashi et al., 2000; Wang et al., 2010). LCK and ZAP70 are translocated to the immunological synapse by RhoH (negative rank 2) (Chae et al., 2010). The ZAP70 target, LCP2 (negative rank 4), is an adaptor protein required for TCR-induced activation and mediates integration of TCR and CD28 co-stimulation signaling by activating VAV1 (negative rank 8), which is required for TCR-induced calcium flux and signal transduction (Dennehy et al., 2007; Raab et al., 1997; Tybulewicz, 2005). LAT (negative rank 38) is another ZAP70 target, which upon phosphorylation recruits multiple key adaptor proteins for signaling downstream of TCR engagement (Bartelt and Houtman, 2013).

Genes that negatively regulate T cell proliferation have therapeutic potential to boost T cell function. Many of the negative regulators are less well-annotated in the canonical TCR signaling pathway, although functions have been assigned to some. Diacylglycerol (DAG) kinases, DGKA (rank 17) and DGKZ (rank 1) – negative regulators of DAG-mediated signals – were both found to restrain human T cell proliferation following stimulation (Arranz-Nicolás et al., 2018; Chen et al., 2016; Gharbi et al., 2011). The E3 ubiquitin-protein ligase, CBLB (rank 4) and its interacting partner, CD5 (rank 12), work together to inhibit TCR activation via ubiquitination leading to degradation of the TCR (Voisinne et al., 2016). TCEB2 (rank 5) complexes with RNF7 (rank 34), CUL5 (rank 162), and SOCS1 (rank 3), which is a key suppressor of JAK/STAT signaling in activated T cells (Kamura et al., 1998; Liau et al., 2018). UBASH3A (rank 10), TNFAIP3, and its partner TNIP1 (rank 13 and 24, respectively) inhibit TCR-induced NFkB signaling, a critical survival and growth signal for activated CD8^+^ T cells (Düwel et al., 2009; Ge et al., 2017). In addition to these key complexes, genes encoding other less well-characterized cell surface receptors, cytosolic signaling components, and nuclear factors were found to inhibit proliferation (**Figure 2F**), revealing a promising resource set of candidate targets to boost the effects of T cell stimulation.

### Arrayed Delivery of Cas9 RNPs Reveals that Hits Alter Stimulation Responses

We next confirmed the ability of high scoring genes to boost T cell activation and proliferation with arrayed electroporation of individual Cas9 ribonucleoproteins (RNPs) (Hultquist et al., 2016; Schumann et al., 2015). We focused our validation primarily on a set of highly-ranked negative regulators of proliferation due to their therapeutic potential to enhance T cell function when targeted. We first tested the effects of these arrayed gene knockouts on T cell proliferation following TCR stimulation. Briefly, CD8^+^ T cells from four human blood donors were stimulated, electroporated with RNPs, rested for 10 days, labelled with CFSE, and then re-stimulated (**Figure 3A** and **Experimental Procedures**). High-throughput flow cytometry determined proliferation responses in edited and control cells based on CFSE dilutions. This validated the ability of many of the tested gene targets to increase T cell proliferation post stimulation, consistent with their robust effects in the pooled screens (**Figure S3A**). For example, CBLB and CD5 knockout cells showed a marked increase in number of divisions post stimulation compared to controls, persistent across different guide RNAs and blood donors (**Figure 3B**). To systematically quantify cell proliferation across samples, we fitted the CFSE distributions using a mathematical model (Munson, 2010) (**Figure S3B**). This analysis revealed that perturbation of multiple negative regulators of T cell stimulation increased the proliferation index score compared to controls (7 out 10 gene perturbations of negative regulators shown here). *UBASH3A, CBLB, CD5*, and *RASA2* knockout T cells all showed greater than 2-fold increase in the proliferation index compared to non-targeting control cells (**Figure 3C**). Notably, targeting these genes did not increase proliferation in the unstimulated cells, indicating that they are not general regulators of proliferation but instead appear to modulate proliferation induced by TCR signaling. In contrast, guides against gene targets that were depleted in the proliferating cells in the pooled screens (CD3D and LCP2), showed a reduction in the proliferation index compared to the non-targeting controls. This orthogonal gene targeting system demonstrated that the majority of top genes identified by our screens robustly modulate stimulation-dependent proliferation in human CD8^+^ T cells.

**Figure 3.**
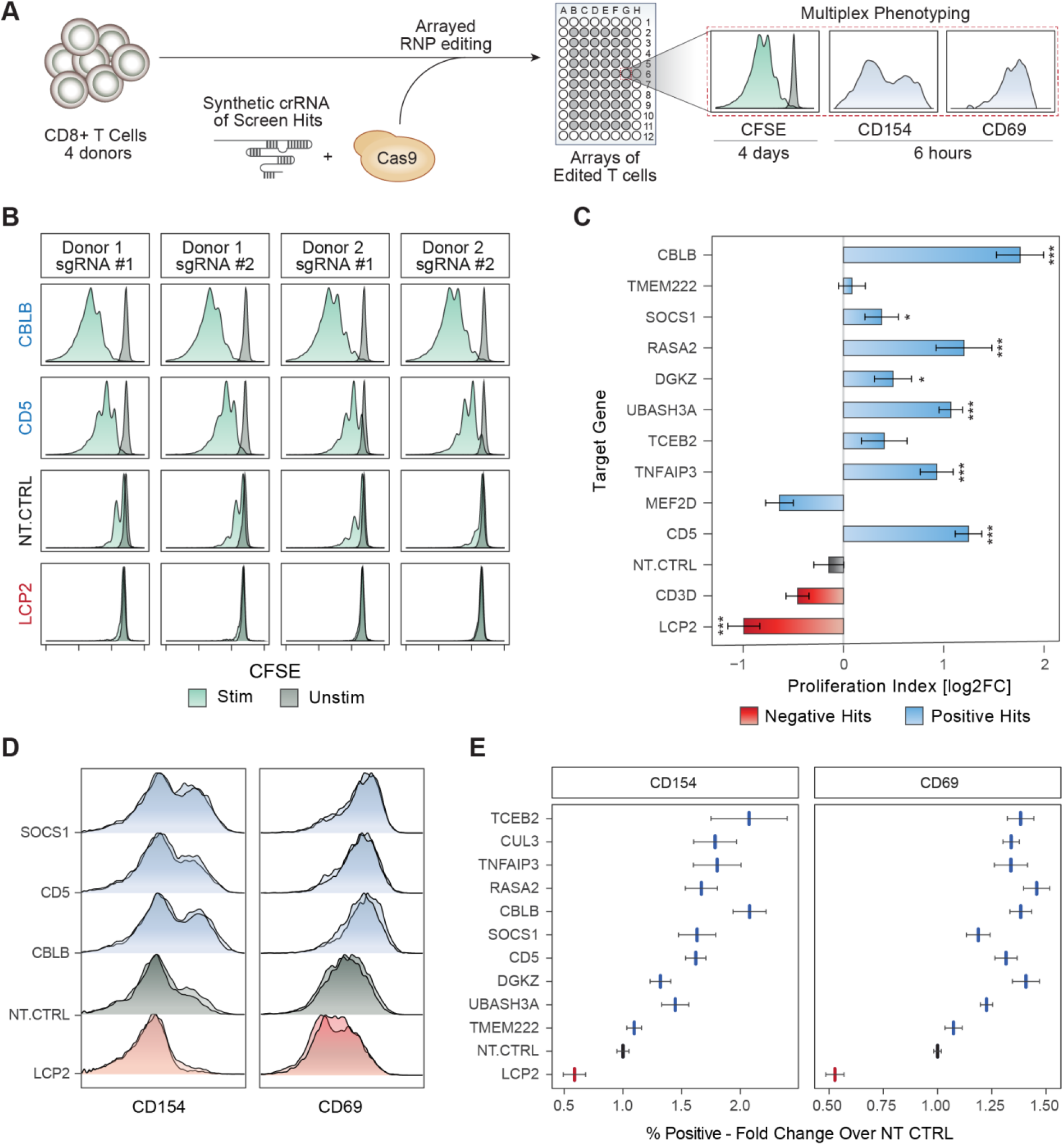
Validation of Gene Targets That Regulate T Cell Stimulation Using RNP Arrays. (A) Overview of orthogonal validation strategy using arrayed electroporation of Cas9 RNPs. (B) Proliferation assay with CFSE-stained CD8^+^ T cells. Each panel shows CFSE signal from TCR-stimulated (green) or unstimulated (dark grey) human CD8^+^ T cells. Shown are data for two guides targeting negative regulators, *CBLB* and *CD5*, compared to non-targeting control (NT-CTRL) guides and guides targeting a critical TCR signaling gene, *LCP2*. (C) Summary of data in (B) across sgRNAs. Gene targets (y-axis) are ordered by their rank in the GW pooled screens. X-axis is the calculated proliferation index (Experimental Procedures), relative to NT-CTRL in each donor (log2 transformed). Bars show mean of two independent experiments, with two donors in each experiment. Error bars are SEM. *** denotes p < 0.001, * denotes p < 0.05, Standard t-test. (D) Early activation markers, as measured by flow cytometry 6 hours post stimulation. Shown are representative distributions of two guides per targeted gene (y-axis) for CD154 (left) and CD69 (right). (E) Summary of data in (D) for all gene targets tested (y-axis). X-axis is the fold-change increase in the marker-positive (CD69 / CD154) population over NT-CTRL. Vertical lines are mean values, error bars are SEM, two guides per gene, for four donors. See also Figure S3.

We next examined whether these hits modulate canonical responses to TCR stimulation in addition to cell proliferation. The phenotype of cells edited in an arrayed format could be assessed with multiple markers at different time points. We analyzed two different cell surface markers of early CD8^+^ T cell activation, CD69 and CD154 (López-Cabrera et al., 1993; Shipkova and Wieland, 2012). At day 10 post-electroporation, cells were assessed 6 hours after re-stimulation. We found that T cells engineered to lack negative regulators of proliferation, such as *SOCS1, CBLB, CD5*, and others, also showed increased surface expression levels of both CD69 and CD154 compared to non-targeting control cells (**Figure 3D** and **Figure S3C-D**). Conversely, targeting a positive regulator of TCR signaling, *LCP2*, reduced expression of CD69 and CD154 in stimulated cells. Overall, the percent of cells expressing these activation markers in each condition was higher for positive hits compared to non-targeting control guides, consistent across two sgRNAs per gene, for four donors (**Figure 3E**). Arrayed editing and phenotyping thus characterized the effects of genetic perturbations and revealed targets that boost stimulation-dependent proliferation and activation programs.

### SLICE Paired with Single Cell RNA-Seq for Molecular Phenotyping of Modified Primary Human Cells

Next, we more deeply characterized the stimulation-dependent transcriptional programs altered by ablation of key target genes in human T cells. The recent combination of pooled CRISPR screens with single-cell RNA-Seq has enabled high-content analysis of transcriptional changes resulting from genetic perturbations in immortalized cell lines (Adamson et al., 2016; Datlinger et al., 2017; Dixit et al., 2016) or cells from transgenic mice (Jaitin et al., 2016). Here, we coupled SLICE with a droplet-based single-cell transcriptome readout for high-dimensional phenotyping of pooled perturbations in primary human T cells. We chose the CROP-Seq platform, as it offers a barcode-free pooled CRISPR screening method with single-cell RNA-Seq using the readily available 10X Genomics platform (Datlinger et al., 2017). We generated a custom library targeting top hits from our GW screen (2 sgRNAs per gene), known checkpoint genes (*PDCD1, TNFRSF9, C10orf54, HAVCR2, LAG3, BTLA*), and 8 non-targeting controls, for a total of 48 sgRNAs (**Table S5**). Human T cells from two donors were transduced with this library, electroporated with Cas9 protein, and enriched with Puromycin selection (**Experimental Procedures**) (**Figure S4A**). Cells were subjected to single-cell transcriptome analysis either with or without re-stimulation to characterize alterations to cell state and stimulation response resulting from each genetic modification.

First, we analyzed the transcriptional states of more than 15,000 resting and stimulated single cells where we could identify an sgRNA barcode. Synthetic bulk gene expression profiles showed that stimulated cells up-regulated many cell cycle genes, indicating response to TCR stimulus (**Figure S4B**). We next visualized the distribution of these single cell transcriptomes in reduced dimensions using Uniform Manifold Approximation and Projection (UMAP) (McInnes and Healy, 2018) (**Figure 4A**). While the unstimulated T cells had donor-dependent basal gene expression patterns, stimulated cells from the two donors tended to have a shared transcriptional signature and clustered together. For example, stimulated cells generally induced expression of cell cycle genes (e.g.: *MKI67*) and cytolytic granzymes (e.g.: *GZMB*) (**Figure 4B**). In contrast, unstimulated cells largely expressed markers of a resting state, such as *IL7R* and *CCR7*. TCR stimulation thus had a strong effect in inducing an activated cell state across biological replicates, although more cells appeared to have been stimulated strongly in Donor 1 than in Donor 2. To systematically impute cell states, we clustered single cells based on their shared nearest neighbors by gene expression (**Figure 4C**). Stimulated cells were enriched in clusters 9-12, which were characterized by preferential expression of mitotic cell cycle and T cell activation cellular programs (**Figure S4C**). This analysis of single cell transcriptomes revealed a characteristic landscape of cell states in human T cells before and after re-stimulation.

**Figure 4.**
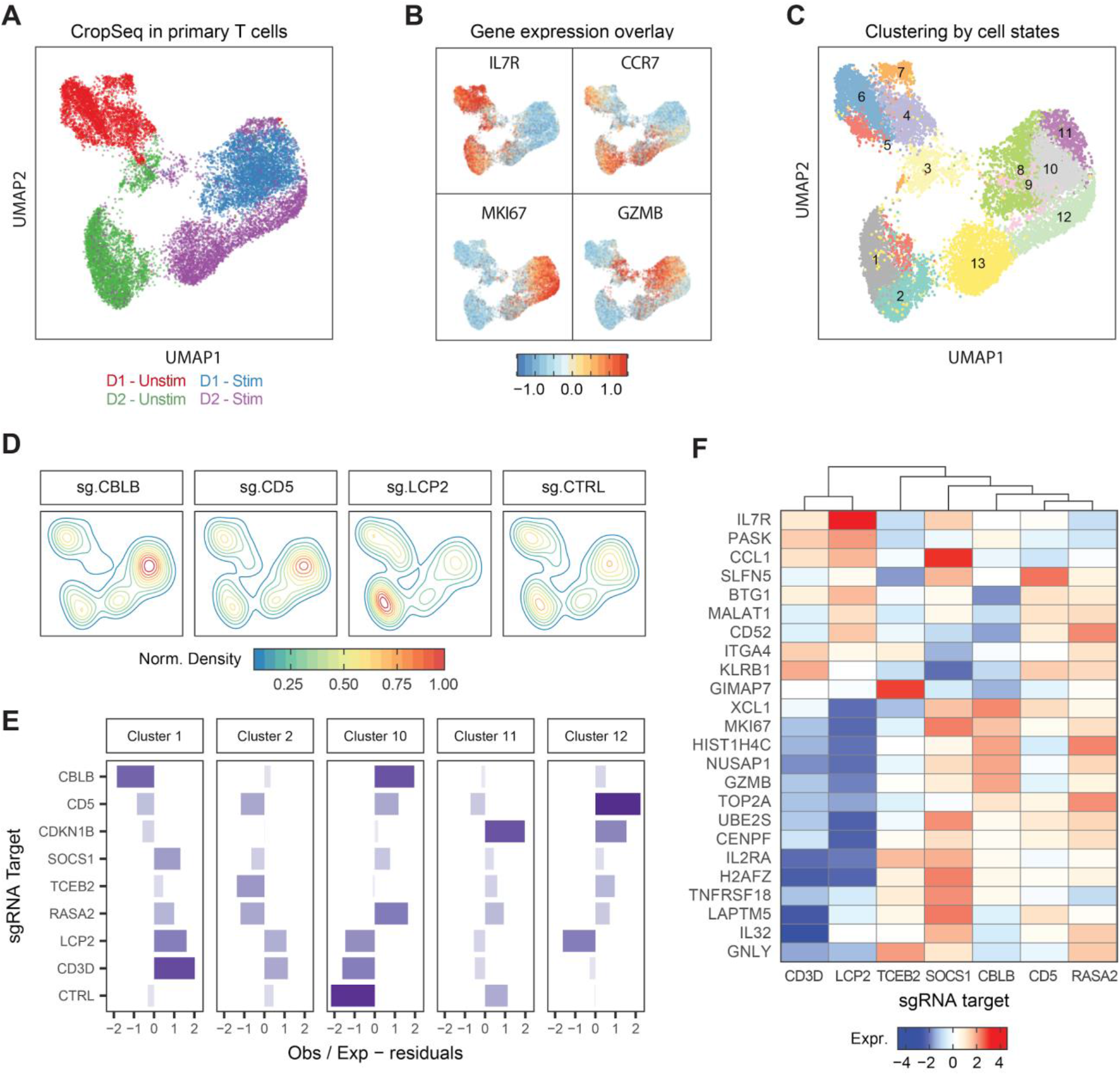
Pairing SLICE with Single Cell RNA-Seq for High Dimensional Molecular Phenotyping of Gene Knockouts in Primary Cells. (A) UMAP plot of all single cells with identified sgRNAs across resting and re-stimulated T cells from two human donors. (B) UMAP with scaled gene expression for four genes showing cluster associations with activation state (IL7R, CCR7), cell cycle (MKI67), and effector function (GZMB). (C) Unsupervised clustering of single cells based on gene expression, 13 clusters identified as labeled. (D) Clustering of cells expressing sgRNAs for CBLB, CD5, LCP and NT-CTRL on the UMAP representation. (E) Y-axis shows over or under representation of cells expressing sgRNAs (y-axis) across clusters (panels), as determined by a chi-square test. (F) Heatmap showing average gene expression (y-axis) across stimulated cells with different sgRNA targets (x-axis). Data is representative of one donor. See also Figure S4.

We next assessed the effect of CRISPR-mediated genetic perturbations on cell states. Efficient editing for the majority of gene targets was validated by reduced expression of sgRNA target transcripts compared to levels in cells with non-targeting control sgRNAs (**Figure S4D**). We tested whether gene perturbations caused altered genetic programs. Cells with non-targeting control sgRNAs were relatively evenly distributed among clusters. In contrast, cells with *CBLB* and *CD5* sgRNAs were enriched in clusters associated with proliferation and activation, and those with *LCP2* sgRNAs were found mostly in clusters characterized by resting profiles (**Figure 4D**). We then quantified which sgRNA targets pushed cells towards distinct cell-state clusters based on their transcriptional profiles (**Figure 4E**). Targeting multiple negative regulators identified in the GW screen such as *CD5, RASA2, SOCS1*, and *CBLB* promoted the cluster 10-12 programs. Perturbation of negative regulators induced markers of activation states (*IL2RA, TNFRSF18/GITR*), cell cycle genes (*MKI67, UBE2S, CENPF* and *TOP2A*), and effector molecules (*GZMB, XCL1*) (**Figure 4F** and **Figure S4E**). In contrast, sgRNAs targeting CD3D or LCP2 inhibited the cluster 10 activation program and promoted the cluster 1-2 programs. SLICE paired with single cell RNA-Seq revealed how target gene manipulation shapes stimulation-dependent cell states.

Targeting different negative regulators of proliferation led to distinct transcriptional consequences. Knockout of *CBLB* tended to induce a cell state signature similar to targeting known checkpoint genes *BTLA* or *LAG3*, as evidenced by similarity in cluster representation (**Figure S4F**). A different shared activation program was observed as a result of targeting *CD5, TCEB2, RASA2* or *CDKN1B*. The integration of SLICE pooled CRISPR screens and single cell RNA-Seq provides a powerful approach to both discover and characterize critical gene pathways in primary human cells. These data also demonstrate that targeting negative regulators of proliferation can induce specialized stimulation-dependent effector gene programs that could enhance the potency of T cells.

### Screen Hits Boost Cancer Cell Killing in vitro by Engineered Human T Cells

Cells engineered to have an enhanced proliferation response and to boost effector gene programs in response to TCR stimulation could hold promise for cancer immunotherapies. We tested the effects of target gene knockout in an antigen-specific *in vitro* cancer cell killing system (**Figure 5A**). Specifically, we used an RFP-expressing A375 melanoma cell line, which expresses the tumor antigen NY-ESO, as a target cell (Wargo et al., 2009). Antigen specific T cells were generated by transduction with the NY-ESO1-reactive 1G4 TCR (Robbins et al., 2008) (**Figure S5A**). These transduced T cells were able to induce caspase-mediated cell death in the target A375 cells, which was exhibited by a rise in the level of caspase and a decline in the level of RFP-tagged A375 nuclei over time (**Figure S5B**). Here, NY-ESO TCR+ T cells were generated from four donors using lentiviral transduction and then edited with RNPs in an array of 24 guides, targeting 11 genes, including non-targeting controls). Antigen-specific T cells with or without gene deletion were then co-cultured with the A375 cells and killing was assessed by quantifying RFP-labeled A375 cells by live time-lapse microscopy over a span of 36 hours.

**Figure 5.**
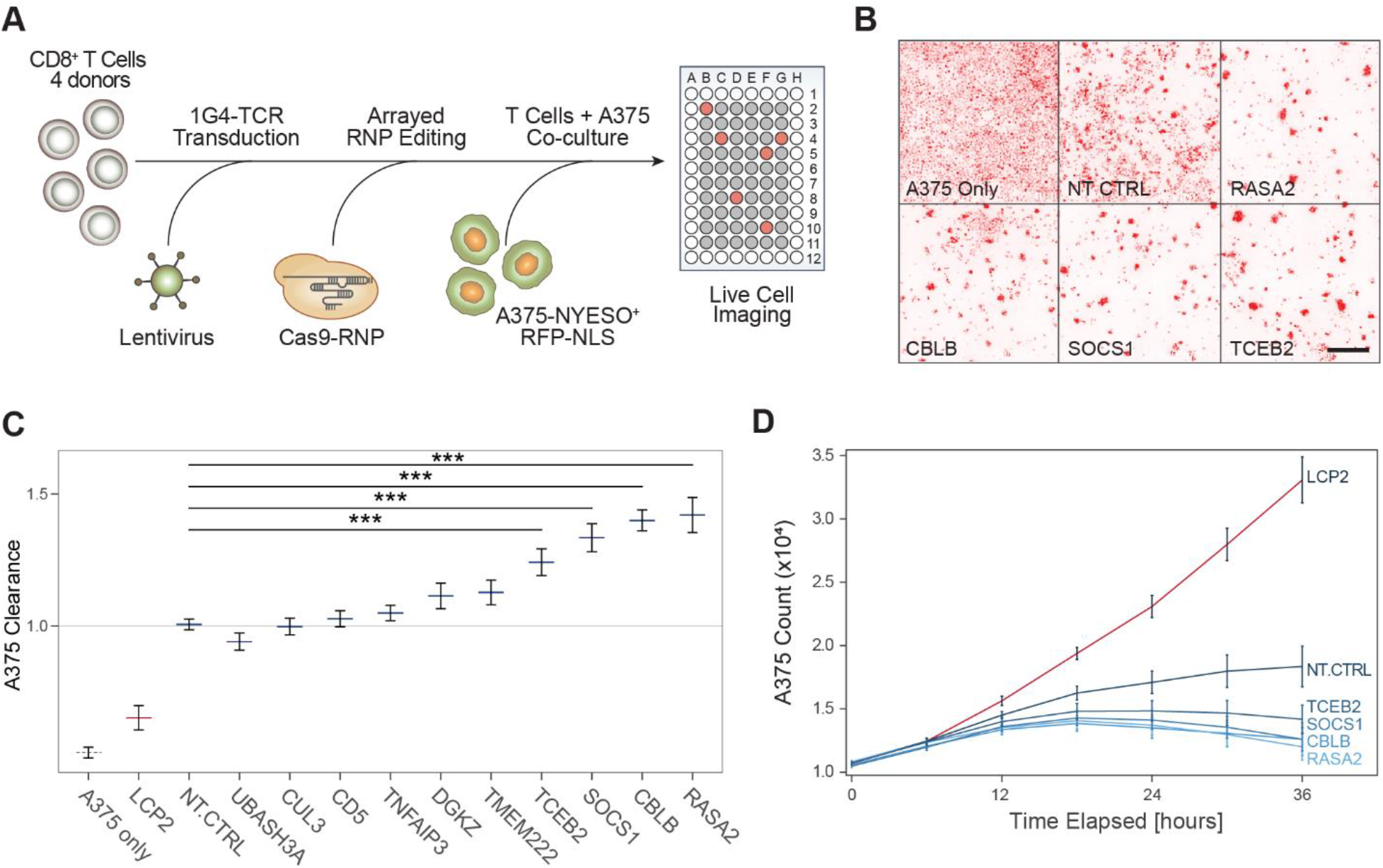
Genome-wide Screen Hits Boost *in Vitro* Cancer Cell Killing by Engineered Antigen-specific Human T Cells. (A) Diagram of a high throughput experimental strategy to test for gene targets that boost cancer cell killing *in vitro* by CD8^+^ T cells. (B) Representative images taken at 36 hours post co-culture of human CD8^+^ T cells and A375-RFP+ tumor cells. Cancer cell density is shown in the red fluorescence channel, for representative wells, as annotated at the bottom left of each panel. Scale bar is 500μm. (C) Clearance of RFP-labeled A375 cells by antigen-specific CD8 T cells after 36 hours. Clearance is defined as count of A375 cells in each well normalized to counts of A375 cells in wells with NT-CTRL T cells. Horizontal lines are the mean, error bars are the SEM, for two guides per gene target, across four donors and two technical replicates. *** denotes p < 0.001, Wilcoxon Rank Sum test. (D) Time traces for A375 cell counts as measured by IncuCyte software for selected hits. Lines are mean for four donors, two guides per target gene. Error bars are SEM. See also Figure S5.

We compared the kinetics of cancer cell killing between gene-edited and control NY-ESO TCR+ T cells. NY-ESO specific T cells started to cluster around RFP+ tumor cells at 12 hours, with more efficient cancer cell clearance at 36 hours for certain sgRNA targets compared to non-targeting controls (**Figure 5B**). As expected, knockout of *LCP2 –* identified in our screens as essential for a strong TCR stimulation response – robustly disabled T cell killing of A375 cells. In contrast, CRISPR-ablation of negative regulators *TCEB2, SOCS1, CBLB* and *RASA2* each significantly increased tumor cell clearance compared to control T cells electroporated with a non-targeting guide RNAs (**Figure 5C**). Targeted deletion of these four genes led to improved kinetics of cancer cell clearance in our assay compared to the non-targeting control conditions (**Figure 5D** and **Figure S5C-D**). Among these, CBLB has been best studied as an intracellular immune checkpoint that can be targeted in T cells to improve tumor control in mouse models (Chiang et al., 2007; Hinterleitner et al., 2012). Targeting *SOCS1*, a negative regulator of JAK/STAT signaling in T cells, showed enhanced T cell clearance comparable to *CBLB* (Liau et al., 2018). Ablation of TCEB2, a binding partner of SOCS1, also improved tumor clearance, suggesting that the SOCS1/TCEB2 complex restrains T cell responses and is a potential target for immunotherapy (Ilangumaran et al., 2017; Kamizono et al., 2001; Liau et al., 2018). *RASA2*, a GTPase-activating protein that stimulates the GTPase activity of wild-type RAS (Arafeh et al., 2015; Maertens and Cichowski, 2014), has not been studied in the context of primary T cells and the immune system, but our findings suggest it may be a modulator of TCR signaling and anti-tumor immunity. In summary, several gene targets identified in the genome-wide screen for proliferative response to stimulation also potentiated *in vitro* tumor killing activity.

### SLICE Screen for Resistance to Immunosuppressive Adenosine Signaling

Adoptive cell therapies that are effective for the treatment of solid tumors will require cells that can respond robustly to tumor antigens even in immunosuppressive tumor microenvironments. With genome editing, T cells could be rendered resistant to particular immunosuppressive cues, and it will be important to identify the relevant T cell pathways for modification. We reasoned that our SLICE screening platform also could be used to identify gene deletions that allow T cells to escape various forms of suppression. We focused on adenosine, a key immunosuppressive factor in the tumor microenvironment (Allard et al., 2017; Beavis et al., 2017; Kazemi et al., 2018). We performed a genome-wide proliferation screen by stimulating T cells in the presence of an adenosine receptor 2 (A2A) agonist (CGS-21680) at a suppressive dose of 20μM (**Figure S6A**) versus a vehicle control for four days (Jacobson and Gao, 2006). We looked for sgRNAs that were enriched in the proliferating cell population (CFSE low) in the A2A agonist treatment condition compared to vehicle (**Figure S6B**, **Table S6**).

While many gene modifications promoted TCR proliferative responses to stimulation in the presence or absence of the adenosine receptor agonist, we identified several sgRNAs that were only enriched in the dividing cells in the presence of CGS-21680 (**Figure 6A**). These gene targets appear to play a selective role in adenosine receptor-mediated T cell suppression. Importantly, *ADORA2A* – encoding the receptor specifically targeted by CGS-21680 – showed a high rank difference between the two treatment conditions (rank 19 in CGS-21680 vs rank 7399 in vehicle control), indicating that its knockout provided a specific escape from CGS-21680 (**Figure 6A** and **Figure S6C**). In contrast, *ADORA2B*, a different adenosine receptor, did not show any proliferative advantage when exposed to this selective A2A agonist, CGS-21680 (**Figure 6A**). These findings encouraged us to investigate other gene targets that show a similar pattern to *ADORA2A* in terms of selective resistance to CGS-21680. We noted that several guanine nucleotide binding proteins with potential roles in adenosine-responsive signaling had higher positive rank scores in the adenosine agonist GW screen, including GCGR (rank 35 vs. 1149), GNG3 (rank 199 vs. 12976), and GNAS (rank 836 vs. 2803). Strikingly, we found multiple guides targeting a previously uncharacterized gene, FAM105A (rank 15 in CGS-21680 vs. rank 13390 in vehicle control), were specifically enriched nearly to the same extent as ADORA2A (**Figure 6B**). Although little is known about FAM105A function, GWAS of allergic diseases implicates a credible missense risk variant in this gene (Ferreira et al., 2017). A neighboring paralogue gene, Otulin (*FAM105B*), encodes a deubiquitinase with an essential role in immune regulation (Damgaard et al., 2016; Fiil and Gyrd-Hansen, 2016) (**Figure S6D**). The screen results suggest a critical role for *FAM105A* in mediating adenosine immunosuppressive signals in T cells.

**Figure 6.**
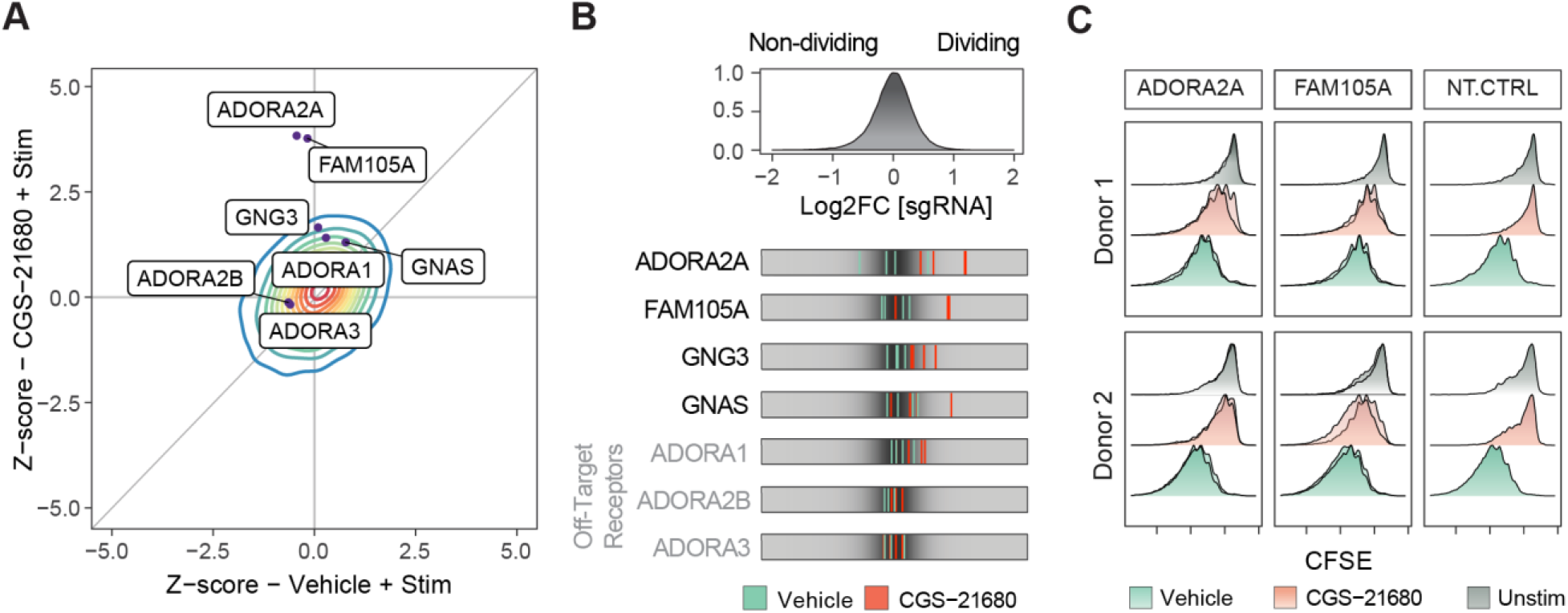
Adapting SLICE to Reveal Resistance to Immunosuppressive Signals in Primary T Cells. (A) Z scores for the genome-wide screen for resistance to adenosine A2A selective agonist CGS-21680 (y-axis) compared to vehicle (x-axis). Contour represents the density (red for higher, blue for lower density) of all genes across the screen. Genes with selective effects on adenosine-mediated immunosuppression deviated upwards from the diagonal identity line. Dots show selected individual gene targets. (B) Top panel: distribution of log2 fold change for all sgRNAs in GW library for T cells treated with CGS-21680 (20μM). Bottom panel: LFC for selected sgRNAs in the vehicle (stimulation only) condition (green) compared to the CGS-21680 treated condition (red). (C) Validation of gene targets from the adenosine resistance screen using T cells edited with individual RNPs. Knockout of both *ADORA2A* and *FAM105A* enables cells to proliferate more robustly in the presence of the adenosine agonist (CGS-21680), compared to the NT-CTRL RNP. Each panel shows results from two independent sgRNAs for two donors. See also Figure S6 and Table S6.

To validate our findings, we used our arrayed RNP platform to edit ADORA2A and FAM105A with CFSE proliferation readout across two donors. We found that targeting each of these genes with two different sgRNAs each led to resistance to suppression by CGS-21680, as predicted by our screen (**Figure 6C**). Importantly, these edits did not lead to increased T cell proliferation in the absence of TCR stimulation, suggesting they selectively overcome CGS-21680 suppression of TCR stimulation. Taken together, SLICE is able to identify both known and novel components of a pathway required for primary cell-response to a specific extracellular cue. Here we identified both extracellular and intracellular targets that could be modified to generate T cells resistant to adenosine suppression. This exemplifies the potential for this platform to be used to discover gene targets to enhance specialized T cell functions.

## DISCUSSION

SLICE provides a new platform for genome-wide CRISPR loss-of-function screens in primary human T cells, a cell type that has been central to a revolution in cancer immunotherapies. SLICE screens can be performed routinely at large-scale in primary cells from multiple human donors, ensuring biologically reproducible discoveries. Here, we enriched for genetic perturbations that enhanced stimulation-responsive T cell proliferation. We elected to use proliferation as our screening output, as it is a broad phenotype regulated by complex genetics. Arrayed editing with Cas9 RNPs allowed us to further characterize the effects of individual perturbations with multiplexed proteomics measured by flow cytometry. Finally, coupling SLICE with single-cell transcriptomics enabled a more global assessment of the functional consequences of perturbing hits from the genome wide screen. Integration of these CRISPR-based functional genetic studies rapidly pinpointed genes in human T cells that can be targeted to enhance stimulation-dependent proliferation, activation responses, effector programs, and *in vitro* cancer cell killing.

The potential power of human primary T cell loss-of-function screens was recently demonstrated in a patient who received CAR T cell therapy for chronic lymphocytic leukemia (CLL) (Fraietta et al., 2018). Non-targeted integration of lentiviral provirus encoding a CAR construct can randomly disrupt endogenous genes. T cells with a lentiviral integration disrupting *TET2* showed strong preferential expansion at the peak of a patient’s response and likely contributed to his complete response to CAR T cell therapy. This patient happened to have a pre-existing hypomorphic mutation in the second allele of *TET2*, suggesting that in this case, *TET2* knockout was key to the T cells proliferative advantage and tumor control. SLICE now provides an opportunity to search more systematically for genetic perturbations that enhance cell expansion and effector function for adoptive T cell therapies.

We found that ablation of at least four targets (SOCS1, TCEB2, RASA2, and CBLB) in human T cells enhanced proliferation and *in vitro* anti-cancer function. Of these, CBLB has been studied in mouse models as an intracellular checkpoint that can be targeted to enhance tumor control by CD8^+^ T cells. Our data suggest that RASA2 and the SOCS1/TCEB2 complex members may also be fruitful targets for modulation in adoptive T cell cancer immunotherapies. These studies demonstrate the potential of SLICE as a tool in primary human CD8^+^ T cells to rapidly discover and validate relevant candidate targets for developing novel immunotherapies. Looking forward, SLICE pooled screens could be adapted to select for perturbations that confer even more complex phenotypes on human T cells, including *in vivo* functions relevant for T cell therapies.

The SLICE pooled screening approach is flexible and versatile as it can be adapted to probe diverse genetic programs that regulate primary T cell biology. Primary T cell screens can be performed with various extracellular selective pressures and/or FACS-based phenotypic selections. We focused on CD8^+^ T cells, but showed that SLICE also can be employed in CD4^+^ T cells, and may be generalizable to many other primary cell types. We demonstrated that a suppressive pressure can be added to the screen, in our case an adenosine agonist, to identify gene perturbations that confer resistance. Future screens could be designed to overcome additional critical suppressive forces in the tumor microenvironment, such as suppressive cytokines, metabolites, nutrient depletion, or suppressive cell types including regulatory T cells or myeloid-derived suppressor cells. In summary, we have developed a novel pooled CRISPR screening technology with the potential to explore unmapped genetic interactions in primary human cells and to guide the design of breakthrough cell therapies.

## ACKNOWLEDGEM ENTS

We thank members of the Marson and Ashworth labs as well as Jeffrey Bluestone, Kole Roybal, Chun Jimmie Ye and Art Weiss for helpful suggestions and technical assistance. We thank Jim Wells and Amy Weeks for experimental insights throughout the conception and realization of the project. We thank Tom Norman and Jonathan Weissman for helpful discussions on Crop-Seq data, Chris Jeans (Macrolab) for reagents, Vin Nguyen and Eunice Wan for technical assistance with flow cytometry and single cell RNA-seq. The 1G4-TCR vector was a gift from Cristina Puig-Saus and Antoni Ribas. This research was supported by grants from the Parker Institute for Cancer Immunotherapy (PICI) and gifts from Jake Aronov and Galen Hoskin (A.M.). A.M. holds a Career Award for Medical Scientists from the Burroughs Wellcome Fund and is an investigator at the Chan Zuckerberg Biohub. J.C. is a Damon Runyon fellow. The UCSF Flow Cytometry Core was supported by the Diabetes Research Center grant NIH P30 DK063720.

## AUTHOR CONTRIBUTIONS

Conceptualization: E.S., J.C., A.A. and A.M., Methodology: E.S., J.C., Investigation: E.S., J.C., V.T., T.L.R, J.M.W., Resources: C.B. and J.L., Formal analysis: E.S., J.C., Software: E.S., Data Curation: E.S., J.C., Supervision: A.A., A.M., Funding acquisition: A.A., A.M., Writing – original draft preparation: E.S., J.C., A.A., A.M., Writing – review and editing: E.S., J.C., V.T., T.L.R, J.M.W., C.B., J.L., M.D., A.A., A.M.

## SUPPLEMENTAL TABLES

**Table S1. Synthetic oligos used in this study: sgRNA library for targeting cell surface and TCR signaling genes, primers and synthetic crRNA for RNP arrays.**

**Table S2. Output of MAGeCK analysis for pilot targeted screen, normalized counts and results for sgRNA and gene level enrichment.**

**Table S3. Output of MAGeCK analysis for primary genome-wide screen.**

**Table S4. Output of MAGeCK analysis for integrated replicates genome-wide screens. Table S5. Oligo sequences for Crop-Seq library.**

**Table S6. Output of MAGeCK analysis for CGS-21680 and vehicle genome-wide screens.**

All supplemental tables are available on request.

## EXPERIMENTAL PROCEDURES

### Isolation and Culture of Human CD8+ T Cells

Primary human T cells for all experiments were sourced from one of two origins: (1) residuals from leukoreduction chambers after Trima Apheresis (Blood Centers of the Pacific) or (2) fresh whole blood samples under a protocol approved by the UCSF Committee on Human Research (CHR#13-11950). Peripheral blood mononuclear cells (PBMCs) were isolated from samples by Lymphoprep centrifugation (STEMCELL, Cat #07861) using SepMate tubes (STEMCELL, Cat #85460). CD8^+^ T cells were isolated from PBMCs by magnetic negative selection using the EasySep Human CD8^+^ T Cell Isolation Kit (STEMCELL, Cat #17953) and used directly. When frozen cells were used (IncuCyte experiments), previously isolated PBMCs that had been frozen in Bambanker freezing media (Bulldog Bio, Cat #BB01) were thawed, CD8^+^ T cells were isolated using the EasySep isolation kit previously described, and cells were rested in media without stimulation for one day prior to stimulation. Cells were cultured in X-Vivo media, consisting of X-Vivo15 medium (Lonza, Cat #04-418Q) with 5% Fetal Calf Serum, 50mM 2-mercaptoethanol, and 10mM N-Acetyl L-Cysteine. After isolation, cells were stimulated with plate-bound anti-human CD3 (Cat #40-0038, clone UCHT1) at 10μg/mL and anti-human CD28 (clone CD28.2) at 5μg/mL (Tonbo, Cat #40-0289) with IL-2 at 50U/mL, at 1e6 cells/mL.

### Lentiviral Production

HEK 293T cells were seeded at 18 million cells in 15 cm poly-L-Lysine coated dishes 16 hours prior to transfection and cultured in DMEM + 5% FBS + 1% pen/strep. Cells were transfected with the sgRNA transfer plasmids and 2nd generation lentiviral packaging plasmids, pMD2.G (Addgene, Cat #12259) and psPAX2 (Addgene, Cat #12260) using the lipofectamine 3000 transfection reagent per the manufacturer’s protocol (Cat #L3000001). The following day, media was refreshed with the addition of viral boost reagent at 500x as per the manufacturer’s protocol (Alstem Cat #VB100). The viral supernatant was collected 48 hours post transfection and spun down at 300g for 10 minutes, to remove cell debris. To concentrate the lentiviral particles, Alstem precipitation solution (Alstem Cat #VC100) was added, mixed, and refrigerated at 4°C for four hours. The virus was then concentrated by centrifugation at 1500g for 30 minutes, at 4°C. Finally, each lentiviral pellet was resuspended at 100x of original volume in cold PBS and stored until use at -80°C.

### Lentiviral Transduction and Cas9 Electroporation

24 hours post stimulation, lentivirus was added directly to cultured T cells at a 1:300 v/v ratio and gently mixed by tilting. Following 24 hours, cells were collected, pelleted, and resuspended in Lonza electroporation buffer P3 (Lonza, Cat #V4XP-3032) at 20e6 cells / 100μL. Next, Cas9 protein (MacroLab, Berkeley, 40μM stock) was added to the cell suspension at a 1:10 v/v ratio. Cells were electroporated at 20e6 cells per cuvette using the pulse code EH115 (Lonza, cat #VVPA-1002). The total number of cells for electroporation was scaled as required. Immediately after electroporation, 1mL of pre-warmed media was added to each cuvette and cuvettes were placed at 37 degrees for 20 minutes. Cells were then transferred to culture vessels in X-Vivo media containing 50U/mL IL-2 at 1e6 cells /mL in appropriate tissue culture vessels. Cells were expanded every two days, adding fresh media with IL-2 at 50U/mL and maintaining the cell density at 1e6 cells /mL.

### CFSE Staining

Cultured cells were collected, spun, washed with PBS, and then resuspended at 1-10 million cells/mL in PBS. CFSE (Biolegend, Cat #423801) was prepared per the manufacturer’s protocol to make a 5mM stock solution in DMSO. At time of use, this stock solution was diluted 1:1000 in PBS for a 5μM working solution, and then added in a 1:1 v/v ratio to the cell suspension. After mixing, cells were incubated for 5 minutes in the dark at room temperature. The stain was then quenched with a volume of media that was 5x the stain volume (e.g. 2ml + 10ml), and incubated for one minute at room temperature in the dark. Cells were then spun down and resuspended in X-vivo media prior to restimulation.

For CFSE staining of arrayed cells that were edited with RNPs, cells were collected from multiple replicate plates and combined into a deep well 96-well plate. Cells were spun down in the deep-well plate and after decanting the media, resuspended in 1mL of PBS per well using a manual multichannel pipette. CFSE was prepared to make a 5uM working solution in PBS per the manufacturer’s protocol as described above. Next, 1mL of the 5μM CFSE was then added in a 1:1 v/v ratio to each well of cells using the multichannel pipette. After mixing, cells were incubated for 5 minutes in the dark at room temperature. The stain was then quenched with 2mLs of X-Vivo media using a multichannel pipette, and incubated for one minute at room temperature in the dark. Cells were then spun down in the deep-well plate, CFSE was decanted, and then cells were resuspended in X-Vivo media prior to restimulation.

### T Cell Proliferation Screen Pipeline

PBMCs from multiple healthy human donors were isolated from TRIMA residuals, as above. After CD8^+^ T cells isolation (Day 0), cells were stimulated with plate-bound anti-human CD3/CD28 and IL-2 at 50U/mL. The following day, 24 hours after stimulation (Day 1), cells were transduced with concentrated lentivirus encoding the pooled sgRNA library, as above. 24 hours after transduction (Day 2), cells were electroporated with Cas9 protein. Cells were then cultured in media with IL-2 at 50U/mL and split every two days, keeping a density of 1e6 cells/mL. On day 14, cells were CFSE stained and then restimulated with ImmunoCult Human CD3/CD28/CD2 T Cell Activator (STEMCELL, Cat #10970). ImmunoCult was used at 1/16 of the dose recommended by the manufacturer, which is 25μL/1e6. Four days later cells were FACS sorted based on CFSE level. Specifically, we defined the non-proliferating cells as those with the highest CFSE peak, and the highly proliferative cells as in the 3rd highest CFSE peak and below (**Figure S1G**). For the T cell proliferation screen with adenosine receptor 2A agonist, CGS-21680 (TOCRIS, Cat #1063), CGS-21680 was first resuspended in DMSO for a stock solution of 10mM, and then added to media for a final concentration of 20μM.

### Preparation of gDNA for Next-Generation Sequencing

After cell sorting and collection, genomic DNA was isolated from cell pellets using a genomic DNA isolation kit (Machery-Nagel, Cat #740954.20). Amplification and bar-coding of sgRNAs for the cell surface sublibrary was performed as described by Gilbert et al. (Gilbert et al., 2014). For the genome-wide screen, after gDNA isolation, sgRNAs were amplified and barcoded as in Joung et al. (Joung et al., 2017), with adaptation to using a two-step PCR protocol. Each sample was first divided into multiple 100μL reactions with 4μg of gDNA per reaction. Each reaction consisted of 50μL of NEBNext 2x High Fidelity PCR Master Mix (NEB, cat #M0541L), 4μg of gDNA, 2.5μL each of the 10μM read1-stagger-U6 and TRACR-read2 primers, and water to 100μL total. The PCR cycling conditions were: 3 minutes at 98°C, followed by 10 seconds at 98°C, 10 seconds at 62°C, 25 seconds at 72°C, for 20 cycles; and a final 2 minute extension at 72°C. After the PCR, all reactions were pooled for each sample and then purified using Agencourt AMPure XP SPRI beads (Beckman Coulter, cat #A63880) per the manufacturer’s protocol. Next 5μL was taken from each purified PCR product to go into a second PCR for indexing. Each reaction included 5μL of PCR product, 25μL of NEBNext 2x Master Mix (NEB, cat #M0541L), 1.25μL each of the 10μM p5-i5-read1 and read2-i7-p7 indexing primers, and water to 50μL total per reaction. The PCR cycling conditions for the indexing PCR were: 3 minutes at 98°C, followed by 10 seconds at 98°C, 10 seconds at 62°C, 25 seconds at 72°C, for 10 cycles; and a final 2 minute extension at 72°C. Post PCR, the samples were SPRI purified, quantified using the Qubit ssDNA high sensitivity assay kit (Thermo Fisher Scientific, cat #Q32854), and then analyzed on the 2100 Bioanalyzer Instrument. Samples were then sequenced on a HiSeq 4000 instrument (Illumina).

### Arrayed Cas9 Ribonucleotide Protein (RNP) Preparation and Electroporation

Lyophilized crRNAs and tracrRNAs (Dharmacon) were resuspended in 10 mM Tris-HCL (7.4 pH) with 150 mM KCl at a stock concentration of 160 μM and stored in -80°C until use. To prepare Cas9-RNPs, crRNAs and tracrRNAs were first thawed, mixed at a 1:1 v/v ratio, and incubated at 37°C for 30 minutes to complex the gRNAs. Cas9 protein (Stock 40μM) was added at a 1:1 v/v ratio and incubated at 37°C for 15 min. Assembled RNPs were dispensed into a 96W V-bottom plate at 3μL per well. Cells were spun down, resuspended in Lonza P3 buffer at 1e6 cells per 20μL, and added to a V-bottom plate with RNPs. The cells/RNP mixture was transferred to a 96 well electroporation cuvette plate (Lonza, cat #VVPA-1002) for nucleofection using the pulse code EH115. Immediately after electroporation, 80μL of pre-warmed media was added to each well and incubated at 37°C for 20 minutes. Cells were then transferred to culture vessels with 50U/mL IL-2 at 1e6 cells /mL in appropriate tissue culture vessels.

### Flow Cytometry for the Arrayed Validation

All array-based validation studies were processed in 96-well round-bottom plates and read on an Attune NxT Flow Cytometer with a 96-well plate-reader. For the RNP-based proliferation validation assays for top targets from the genome-wide screen, cells were stained with CFSE in the 96-well format prior to restimulation as described in methods above. For the evaluation of activation marker levels on arrayed RNP-edited cells, the following antibodies were used: CD69 (Biolegend, cat #310904), CD154 (Biolegend, cat #310806), and CD8a (Biolegend, cat #301038).

### Pooled sgRNA Library Construction

For the cloning of the targeted cell surface sublibrary, we followed the custom sgRNA library cloning protocol as described by Joung et al. (Joung et al., 2017). We utilized the pgRNA-humanized backbone (Addgene, plasmid #44248). To optimize this plasmid for cloning the library, we first replaced the sgRNA with a 1.9kb stuffer derived from the lentiGuide-Puro plasmid (Addgene, plasmid #52963) with flanking BfuAI cut sites. This stuffer was excised using the BfuAI restriction enzyme (NEB, #R0701) and the linear backbone was gel purified (Zymo, #D4007). We designed a targeted library to include all genes matching Gene Ontology for “Cell Surface”, “‘T cell receptor signaling pathway”, or “cytokine receptor activity”. In total we included 1211 genes with 4 guides per gene, and 48 non-targeting controls (**Table S1**). Guides were subsetted from the Brunello sgRNA library (Doench et al., 2016), and the pooled oligo library was ordered from Twist Bioscience to match the vector backbone. Oligos were PCR amplified and cloned into the modified pgRNA-humanized backbone by Gibson assembly as described by Joung et al. (Joung et al., 2017). For the genome-wide screens, the Brunello plasmid library in the lentiGuide-Puro backbone (Addgene, cat. 73178) was purchased from Addgene. The library was amplified using Endura ElectroCompetent Cells following the manufacturer’s protocol (Endura, Cat #60242-1).

### SLICE adapted to CROP-Seq

The backbone plasmid used to clone the CROP-Seq library was CROPseq-Guide-Puro, purchased from Addgene (Addgene. Plasmid #86708). This library consisted of 20 gene targets (2 guides per gene selected from hits in the GW screen) and 8 non-targeting control guides, for a total of 48 guides (**Table S5**). Oligos for these library guides were purchased from Integrated DNA Technologies (IDT) and cloned into the CROPseq-Guide-Puro plasmid backbone using the methods described by Datlinger et al. (Datlinger et al., 2017). Lentivirus was produced from this pooled plasmid library and used to transduce CD8^+^ T cells from two healthy donors, as above. 48 hours after transduction, cells were treated with 2.5ug/mL Puromycin for three days, and subsequently sorted for live cells using Ghost Dye 710 (Tonbo Biosciences, cat #13-0871). Two days post sorting, cells were re-stimulated as above. 36 hours post restimulation, cells were collected, counted, and prepared for Illumina sequencing by Chromium™ Single Cell 3’ v2 (PN-120237), as per manufacturer protocol.

### CROP-seq Guide Reamplification

For the guide reamplification, samples were amplified and barcoded using a two-step PCR protocol. First, each sample was divided into 8 PCR reactions with 0.1ng template of cDNA each. Each 25μL reaction consisted of 1.25μL P5 forward primer, 1.25μL Nextera Read 2 reverse primer, priming to the U6 promoter to enrich for guides, 12.5μL NEBNext Ultra II Q5 Master Mix (NEB, cat #M0544L), 0.1ng template, and water to 25μL. The PCR cycling conditions were: 3 minutes at 98°C, followed by 10 seconds at 98°C, 10 seconds at 62°C, 25 seconds at 72°C, for 10 cycles, and a final 2 minute extension at 72°C. After the PCR, all reactions were pooled for each sample and purified using Agencourt AMPure XP SPRI beads per the manufacturer’s protocol. Next, 1μL was taken from each purified PCR product to go into a second PCR for indexing. Each reaction included 1μL of PCR product, 12.5 μL NEBNext Ultra II Q5 Master Mix (NEB, cat #M0544L), 1.25μL P5 forward primer, 1.25μL Illumina i7 primer, and water to 25μL. The PCR cycling conditions were: 3 minutes at 98°C, followed by 10 seconds at 98°C, 10 seconds at 62°C, 25 seconds at 72°C, for 10 cycles, and a final 2 minute extension at 72°C. After the PCR, all reactions were SPRI purified and quantified using the Qubit dsDNA high sensitivity assay kit (Thermo Fisher Scientific, cat# Q32854) and run on a gel to confirm size. Samples were then sequenced on a MiniSeq instrument (Illumina).

### A375 and T cell *in vitro* Co-culture Assay

A375 melanoma cells were transduced with lentivirus to establish an RFP-nuclear tag (IncuCyte, Cat #4478) for optimal imaging on the IncuCyte live cell imaging system. 24 hours after stimulation, CD8^+^ T cells from healthy donors were transduced with virus containing the 1G4 NY-ESO1-reactive α95:LY TCR construct. Five days after transduction, cells were FACS sorted for a pure population of cells expressing the construct using the HLA-A2+ restricted NY-ESO-1 peptide (SLLMWITQC) dextramer-PE (Immundex, cat #WB2696). Cells were then expanded in X-Vivo media containing IL-2 at 50U/mL for a total of 14 days after initial stimulation. For initial optimization of this system, A375 cells were seeded at 24 thousand cells per well and T cells from two donors transduced with the 1G4 NY-ESO specific TCR were added at varying T cell to tumor cell ratios. The IncuCyte Caspase-3/7 red apoptosis reagent (IncuCyte, Cat #4704) was added to each well per the manufacturer’s instructions, and then imaged every 4 hours on the IncuCyte live cell imaging system. In parallel, A375 cells with the RFP-nuclear tag were seeded at 4,000 cells per well, and the same T cells from two donors transduced with the 1G4 TCR were added at the same ratios as in the caspase experiment, and these were imaged in parallel.

To test candidate gene targets, sorted 1G4+ T cells were edited as in the arrayed RNP experiments at day 10 post stimulation. On the day prior to co-culture, A375 cells were seeded at 5,000 cells per well in a 96W plate in 100μL of complete RPMI media. Complete RPMI media includes RPMI (Gibco, cat #11875093), 10% Fetal Calf Serum, 1% L-glutamine, 1% NEAA, 1% HEPES, 1% pen/strep, 50mM 2-mercaptoethanol, and 10mM N-Acetyl L-Cysteine. The next day, 1G4+ edited T cells were added to each well on top of the 5,000 A375 cells at indicated T cell to cancer cell ratios. T cells were added in 50μL of complete RPMI with IL-2 and glucose at final concentrations of 50U/mL and 2g/dL, respectively. Plates were then imaged using the IncuCyte live cell imaging system, where the number of A375 RFP-positive nuclei were counted over time.

### Oligo list

All guide RNAs and primers used in the study are listed in Supplementary Table S1.

## QUANTIFICATION AND STATISTICAL ANALYSIS

### Analysis of Pooled CRISPR Screens

To identify negative and positive hits in our screens, we used the MAGeCK software to quantify and test for guide enrichment (Li et al., 2014). Abundance of guides was first determined by using the MAGeCK “count” module for the raw fastq files. For the genome-wide Brunello libraries, the 5’ trim length was set to remove the staggered offset introduced by the library preparation, by using the parameter: “—trim-5 23,24,25,26,28,29,30”. For the targeted libraries the constant 5’ trim was automatically detected by MAGeCK. We removed guides with an absolute count under 50 in more than 80% of the samples. To test for robust guide and gene-level enrichment, the MAGeCK “test” module was used with default parameters. This step includes median ratio normalization to account for varying read depths. We used the non-targeting control guides to estimate the size factor for normalization, as well as to build the mean-variance model for null distribution, which is used to find significant guide enrichment. All donor replicates in each screen were grouped for analysis to account for biological noise. MAGeCK produced guide-level enrichment scores for each direction (i.e. positive and negative) which were then used for alpha-robust rank aggregation (RRA) to obtain gene-level scores. The p-value for each gene is determined by a permutation test, randomizing guide assignments and adjusted for false discovery rates by the Benjamini–Hochberg method. Log2 fold change (LFC) is also calculated for each gene, defined throughout as the median LFC for all guides per gene target. Where indicated, LFC was normalized to have a mean of 0 and standard deviation of 1 to obtain the LFC Z-score.

### Gene Set Enrichment Analysis for Screen Hits

To find enriched annotations within screen hits, we used Gene Set Enrichment Analysis, as implemented in the fgsea R package (Sergushichev, 2016). The input for enrichment consisted of the LFC values for all genes tested in the screen. We used the KEGG pathways dataset as the reference gene annotation database, including only gene sets with more than 15 members and less than 500 members. For the external gene set for *in vivo* immunotherapy hits shown in Figure 2E, we used the 43 genes as determined by Zhou et al. (Zhou et al., 2014), having 3 or more shRNA guides with over 4-fold enrichment in T cells from tumor tissue compared to spleen. Normalized enrichment scores and p-values were determined by a permutation test with 10,000 iterations with same size randomized gene sets and adjusted with the FDR method.

### Fitting CFSE Distributions for Arrayed Validation Screens

We used the FlowFit R package to extract quantitative parameters from the CFSE profiles across all samples. As CFSE staining for arrays was done for individual populations of edited cells, the signal peak for the parental population might shift slightly from well to well. To account for this, for each well the stimulated well was compared to an identical unstimulated well, expected to have a single peak at the end of the assay. The FlowFit package implements the Levenberg-Marquadt algorithm to estimate the size and position of the parental population peak. We then used the fitted parameters from the unstimulated wells to fit the CFSE profiles of the corresponding stimulated cells. These CFSE profiles are modeled as Gaussian distributions, with log2 distanced peaks resulting from cell divisions and CFSE dilution. The fitted models were inspected visually, adjusting fitting parameters to minimize deviance from the original CFSE signal. The fitted models were used to calculate the proliferation index (Munson, 2010), defined as the total count of cells at the end of the experiment divided by the calculated original starting number of parent cells. This parameter is robust to variation in the starting CFSE staining intensities.

### Analysis of SLICE Paired with Single-cell RNA-Seq

Pre-processing of the Illumina sequencing results from the 10X Genomics V2 libraries was performed with CellRanger software, version 2.1. This pipeline produces sparse numerical matrices for each sample, with gene-level counts of unique molecular identifiers (UMI) identified for all single cells passing default quality control metrics. These gene expression matrices were processed with Seurat R package (Butler et al., 2018), as described elsewhere (https://satijalab.org/seurat/pbmc3k_tutorial.html). Only cells with more than 500 genes identified were used for downstream analyses. Using Seurat, counts were log normalized, regressing out total UMI counts per cell and percent of mitochondrial genes detected per cell, and scaled to obtain gene level z-scores. We then applied principal component analysis (PCA), using the 1,000 most variable genes across cells. The first 30 PCA components were used to construct a uniform manifold approximation and projection (UMAP) to visualize single cells in a two dimensional plot, as in Figure 4A. Gene expression for single cells as displayed in Figure 4B was calculated as log10(UMI count + 1) and scaled. Clustering in Figure 4C, was performed by the Louvain algorithm on the shared nearest neighbor graph, as implemented by the FindClusters command from the Seurat R package. For synthetic bulk differential gene expression in Figure S4B, UMI counts per gene were summed for all cells with non-targeting control guides in each sample, and the DESeq2 R package was used to determine differentially expressed genes. For gene list enrichment analysis in cell clusters, the REACTOME database (Fabregat et al., 2018) was used as reference to generate Figure S4C.

To associate guides with identified cell barcodes, we processed both fastq files from the 10X libraries and from the re-amplification PCR. The read2 files were matched to the guide library using matchPattern as implemented in the R ShortRead package. The pattern used was the sequence of the U6 promoter preceding the guide sequence appended to the 20bp library guide sequences (e.g. TGGAAAGGACGAAACACCGNNNNNNNNNNNNNNNNNNNN, where N denotes the guide sequence), allowing for 4 mismatches total. The mate Read1 pairs for reads with matched guides were used to determine the cell barcode and UMI assignment. We filtered out reads appearing less than twice and cells with more than one assigned guide. The Chi-square test was used to determine over-representation of cells with guides for the same gene target across cell-state driven clusters. Standardized residuals from the chi-square test were scaled and used to generate Figures 4E and S4F.

### DATA AND SOFTWARE AVAILABILITY

Raw sequencing files for all screens performed, raw files for the single-cell RNA-Seq, supplementary tables, and code used to analyze data and produce figures in this work is available by request.

## SUPPLEMENTAL FIGURES

**Figure S1.**
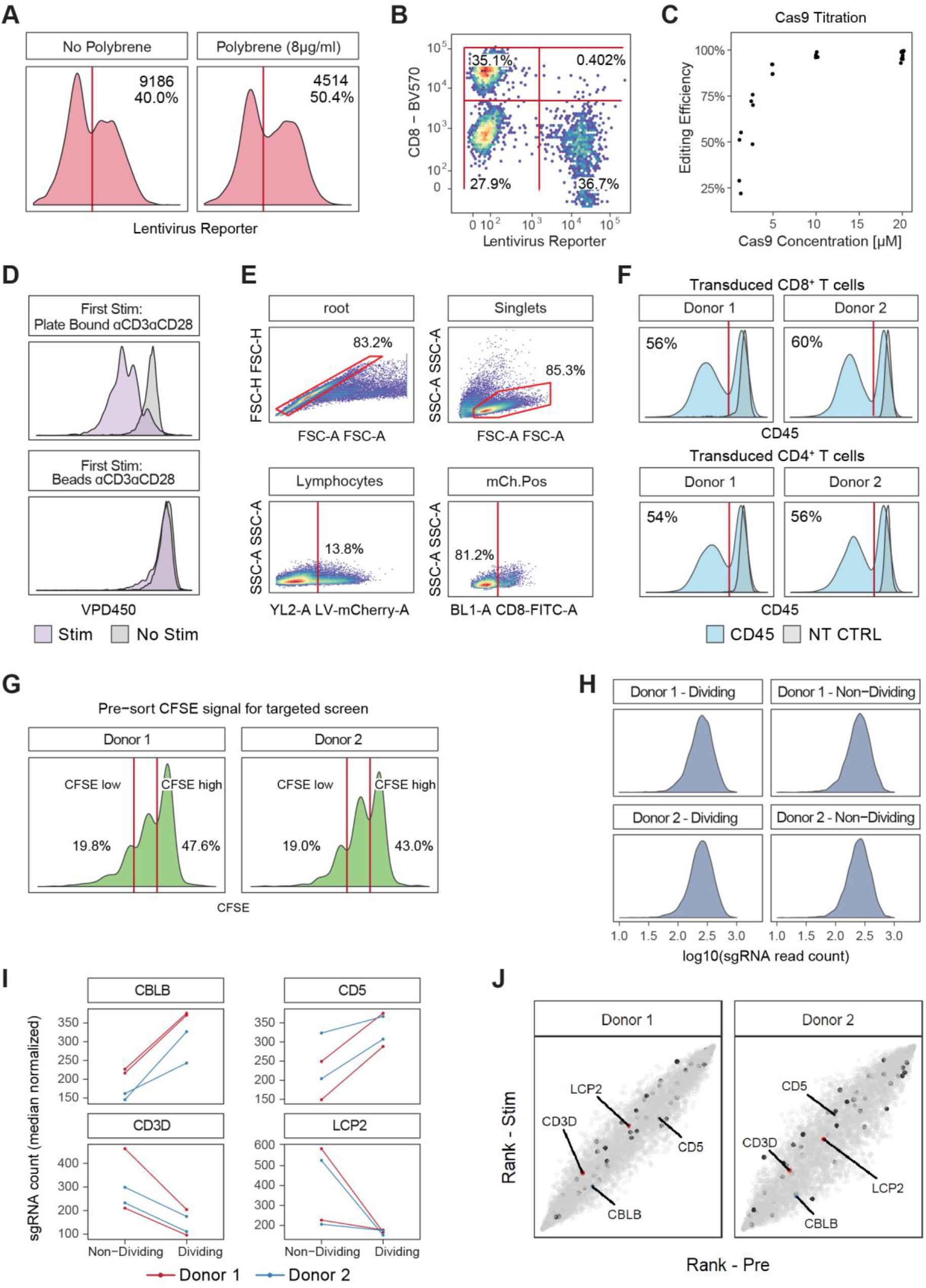
Establishing SLICE in Primary Human T Cells, Related to Figure 1. (A)To optimize SLICE for large scale screens, we tested whether increasing transduction rates with polybrene yielded a higher absolute number of viable edited cells. Although addition of polybrene modestly increased transduction efficiency (bottom labels) - omitting it improved viability and resulted in a higher total number of transduced cells (top labels). (B)FACS plot showing expression of a fluorescent reporter (x-axis) from primary human CD8^+^ T cells transduced with sgRNA lentivirus compared to antibody staining of the targeted protein, CD8a (BV-570, y-axis). Following electroporation of Cas9 protein (3μL in 20μL of cell suspension, stock 20μM), cells expressing the sgRNA were largely CD8 negative. (C)Editing of the CD8a protein (y-axis) for cells from (A), electroporated with different stock concentrations of Cas9 protein. Editing is calculated as the proportion of CD8 negative cells to CD8 positive cells in successfully transduced cells. Editing efficiency depended on Cas9 concentration. Data is shown for two human donors. (D)Effect of first stimulation on response to second stimulation (Day 9) by anti-CD3 and anti-CD28 complexes, as measured by Proliferation dye VPD450. First stimulation with anti-CD3/CD28 beads (bottom panel) at a 1:1 bead to T-cell ratio hindered the ability of CD8^+^ T cells to respond to restimulation. First stimulation using plate-bound antibodies (top panel) maintained the ability of cells to respond, allowing for TCR dependent proliferation screens. (E)Gating strategy for cells shown in Figure 1B. (F)SLICE was effective at knockout of candidate gene target, *CD45*, in both primary human CD8^+^ and CD4^+^ T cells. Non-targeting control sgRNAs (NT-CTRL, grey) showed that knockout is specific to cells transduced with the *CD45* targeting guide (blue). (G)CFSE signal for pre-sort cells in pilot screen as in Figure 1C. (H)Distribution of read counts after deep sequencing of sgRNAs of sorted cell populations, across two donors in the pilot screen. (I)Abundance of top two guides by absolute fold change, for hits from Figure 1C. Each line is a unique sgRNA targeting the given gene, labeled at the top of each panel. Patterns of guide enrichment and depletion are consistent across two guides and two donors. (J) Comparison of rank by log fold change from a parallel screen with the same experimental timeline and the sgRNA library as in Figure 1C, but without CFSE-based enrichment sort. This growth based screen had a low signal to noise ratio, indicated by the proximity to the diagonal for TCR related guides from Figure 1C. Grey dots are individual targeting guides, black dots are non-targeting control (NT-CTRL) guides.

**Figure S2.**
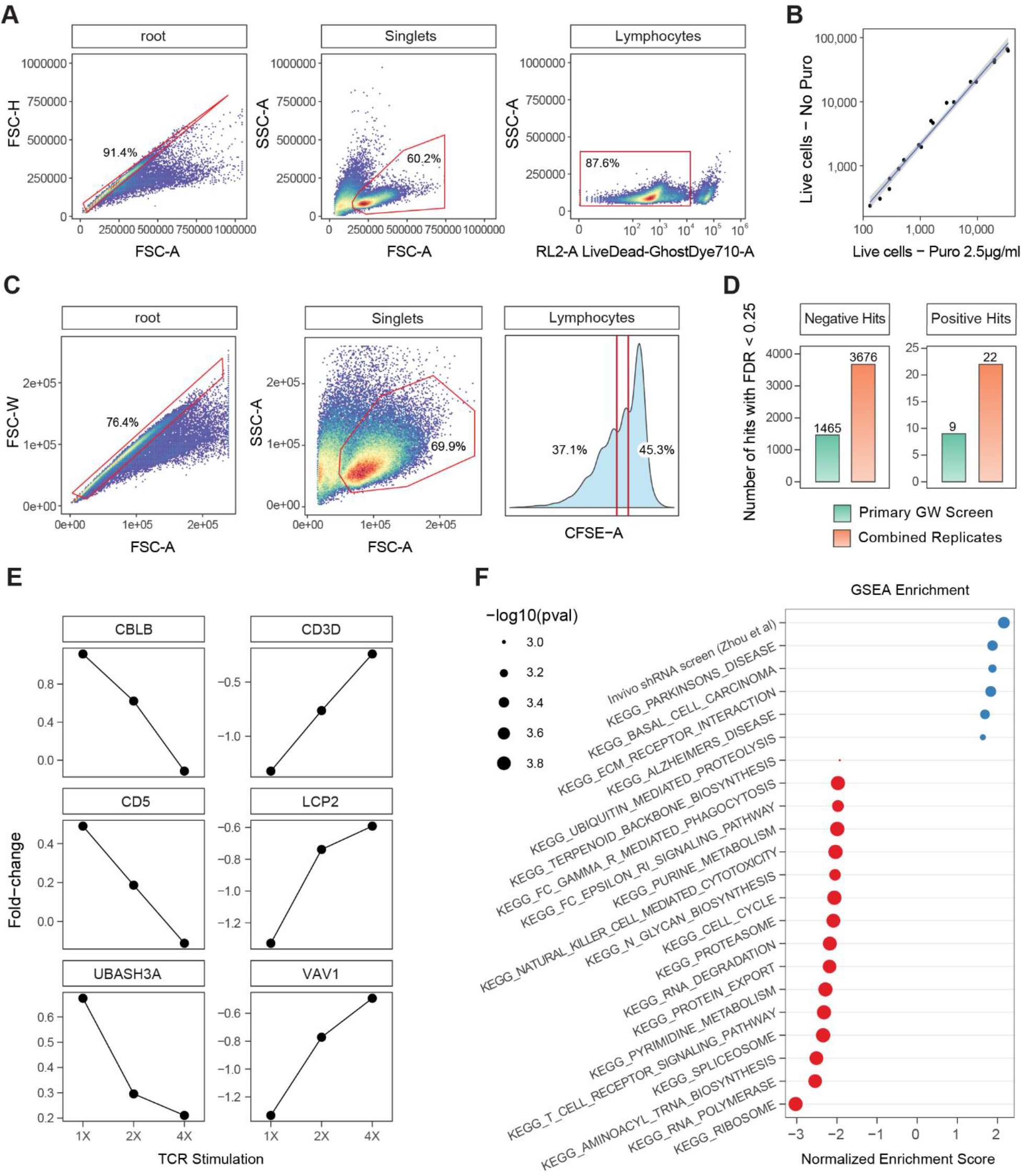
Genome-wide Pooled CRISPR Screens in Primary T cells, Related to Figure 2. (A)Gating strategy for estimating transduction efficiency in the primary GW screen by Puromycin titration. (B)Transduced cells in (A) were cultured for two days with or without Puromycin, in a serial dilution of cell densities. The total number of live cells as in (A) was compared between the two conditions, across cell dilutions. The fitted line is a linear regression, with a slope of 0.51, indicating efficient transduction (R^2^=0.99, shaded area is 95% confidence interval (CI)). (C)Gating and sorting strategy for a representative sample from the GW screen. We sorted cells as indicated in the CFSE panel, allowing for a defined separation between CFSE-high and CFSE-low populations. (D)Comparison of hits discovered in the primary GW screen with two donors (green bars), and the composite data from an independent secondary screen with two more donors (orange bars) - 4 donors in total. Hits were defined by having FDR < 0.25 by the MAGeCK RRA algorithm in each direction. Labels at the top of the bars are the numerical values of the number of hits in each group. (E)Effect of TCR stimulation level (x-axis) on the gene-level fold change for three positive and three negative regulators of T cell stimulation. (F)Enrichment of gene-level fold change by GSEA results for KEGG pathways with FDR < 0.01. In addition, included is a gene set from a published study of shRNAs enriching for T cell tumor infiltration. Each dot is an annotated gene set (y-axis), x-axis shows the normalized enrichment score (NES), compared to mean of randomized gene sets of the same size. Size of each dot is negatively proportional to the p-value of each enrichment.

**Figure S3.**
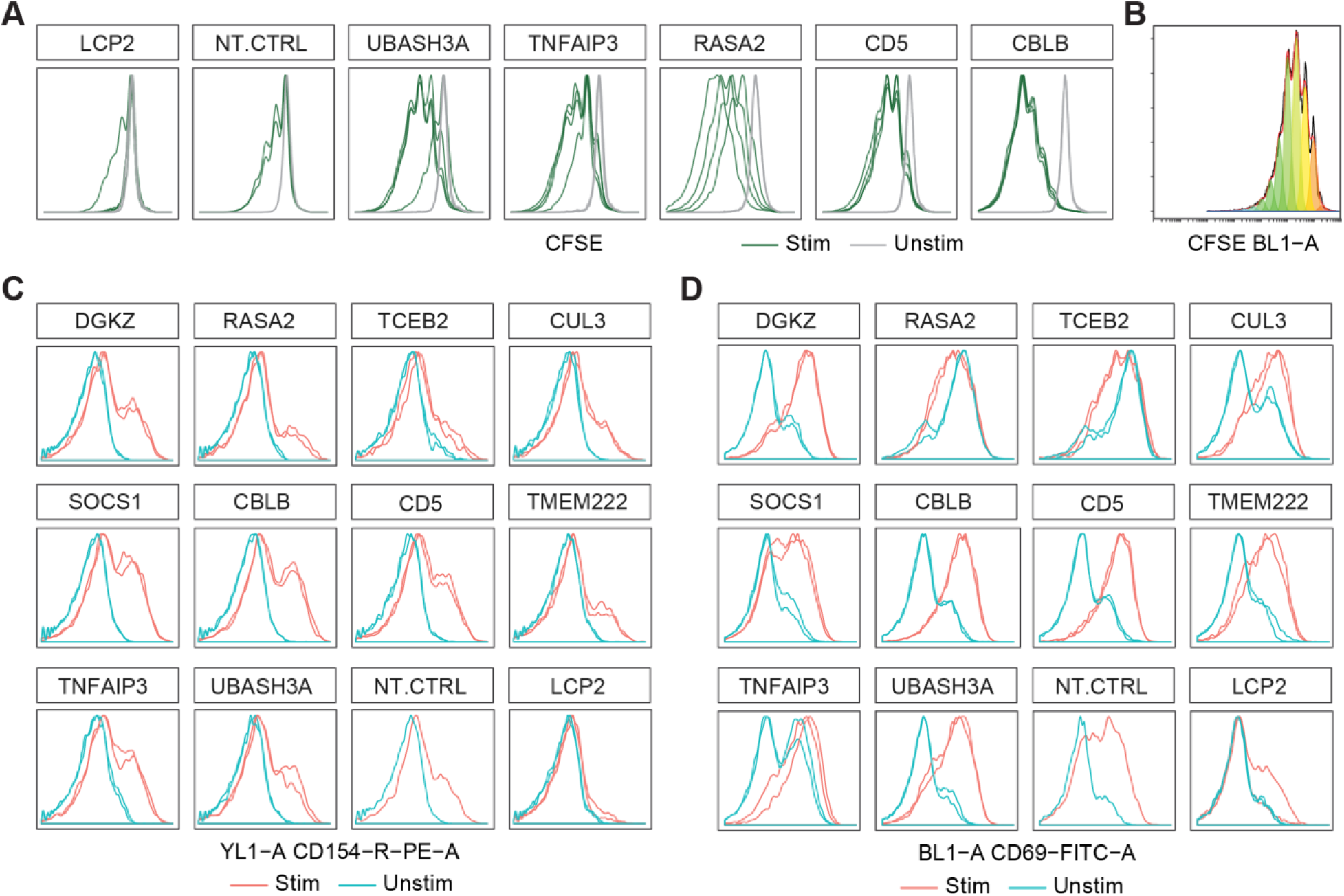
RNP Arrays Validate Hits from Genome-wide Screens, Related to Figure 3. (A)CFSE traces for hits in Figure 3C. Each panel shows two unique guides per gene target, for two technical replicates. Data is representative of one donor. (B)Curve fitting for representative CFSE trace, as in Figure 3C. Panel shows fitted Gaussian distributions for CFSE peaks, as determined by flowFit R package. The extracted parameter is the proliferation index, defined as the total count of cells in all generations divided by the calculated number of original parent cells. (C)Distribution of the activation marker, CD154, across all targets tested. Lines show the measured expression for stimulated (red) and unstimulated (blue) cells, edited with two guides targeting the gene as labeled at top of each panel. Data is representative of one donor. (D)Same as (C), for CD69.

**Figure S4.**
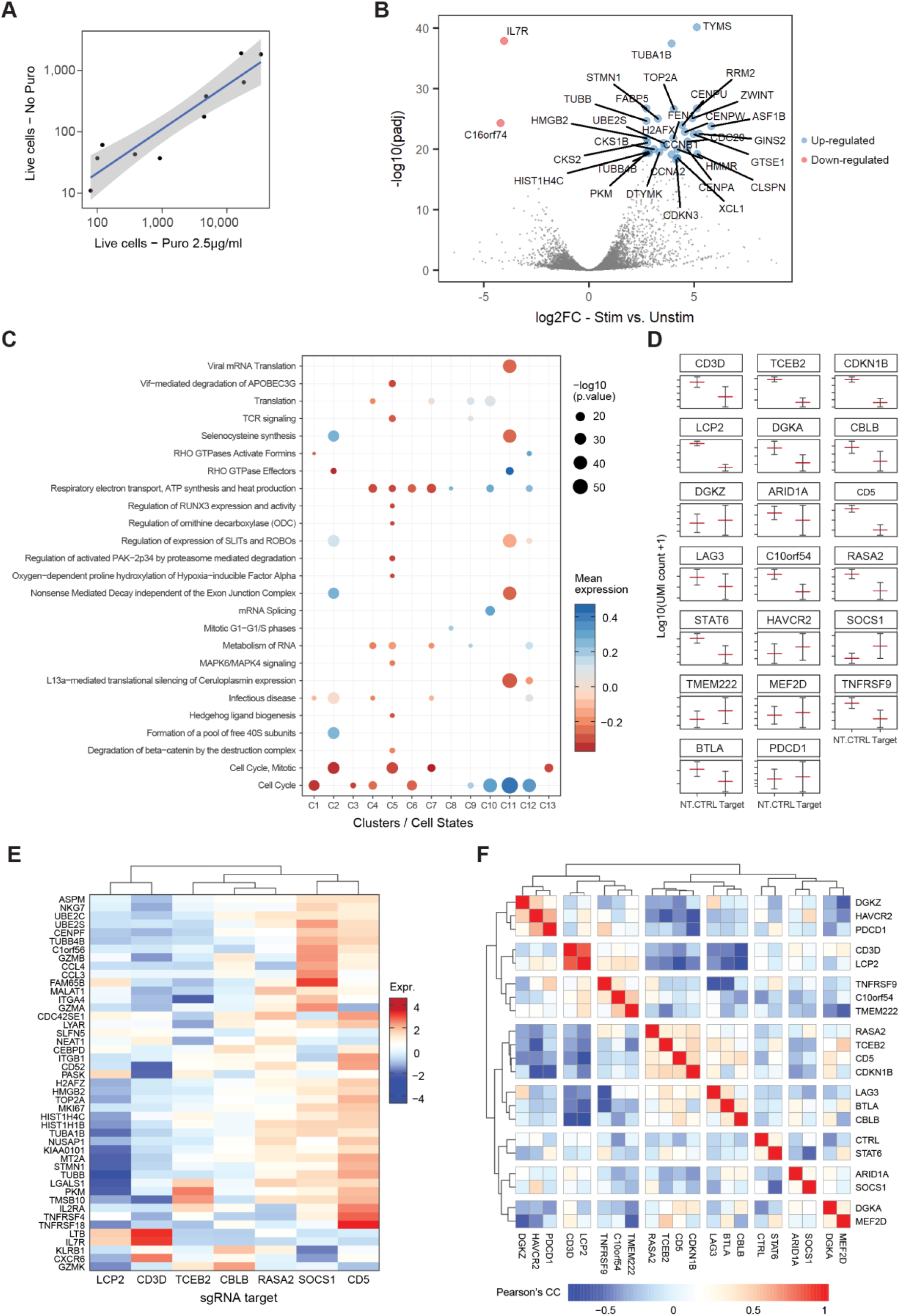
SLICE Coupled with Single-Cell RNA-Seq, Related to Figure 4. (A)Assessment of transduction efficiency for the Crop-Seq experiment by Puromycin selection. We estimated the infection rate to be 13.9% by linear regression (R^2^ = 0.76, blue line, shaded area is 95% CI). This intentionally low transduction rate was used to enrich for cells with only a single integrated sgRNA cassette. (B)Volcano plot for differential gene expression by DESeq2 for synthetic bulk RNA-Seq profiles (collapsed UMI counts per sample) of stimulated cells compared to unstimulated cells. Dots are genes, y-axis shows the significance of enrichment, x-axis shows the magnitude and direction of the log2 fold change (LFC) in transcript abundance. Dots are colored by having |LFC| > 2.5 and adjusted p value < 1e-18, blue denotes genes that are up-regulated in stimulated cells, red denotes genes that are down-regulated. (C)Enrichment of gene annotations in clusters based on single-cell RNA expression profiles. Each dot shows the enrichment of a REACTOME annotation (y-axis) across clusters (x-axis). The genes associated with each cluster were determined by a differential gene expression test, comparing all cells in a cluster to all other cells. The size of each dot is proportional to the significance of the enrichment, color is whether the average expression level of genes in the corresponding annotation is up- or down-regulated (red and blue, respectively) in each cluster. (D)Effects on target gene editing at a single cell level for the CROP-Seq experiment. Each panel shows the mean and standard error of the mean (SEM) for unique molecular identifier (UMI) counts of a targeted gene in the library. This is calculated for each target transcript for all cells either expressing an on-target sgRNA (Target) or a non-targeting control guide (NT.CTRL). Mean is calculated across four samples, two guides per gene. Most gene targets showed a lower level of transcript for cells expressing targeting guides compared to control guides, suggesting successful knockout. We noted that some gene targets were expressed at low levels, thus the effect on target gene editing might be harder to detect at the single cell level. (E)Gene expression profiles for selected sgRNAs, same as Figure 4F, for the second donor. Genes shown are enriched (|logFC| > 1 and adjusted p < 0.05) in clusters 8-12, associated with stimulated cells. Dendrogram is based on Euclidean distance, with the Ward D2 algorithm, as implemented in *hclust* in R. (F)Similarity in cluster association for cells expressing targeting sgRNAs. Shown is the Pearson correlation coefficient for chi-square residuals across all clusters. Dendrogram calculation is as in (E), segmented to three levels.

**Figure S5.**
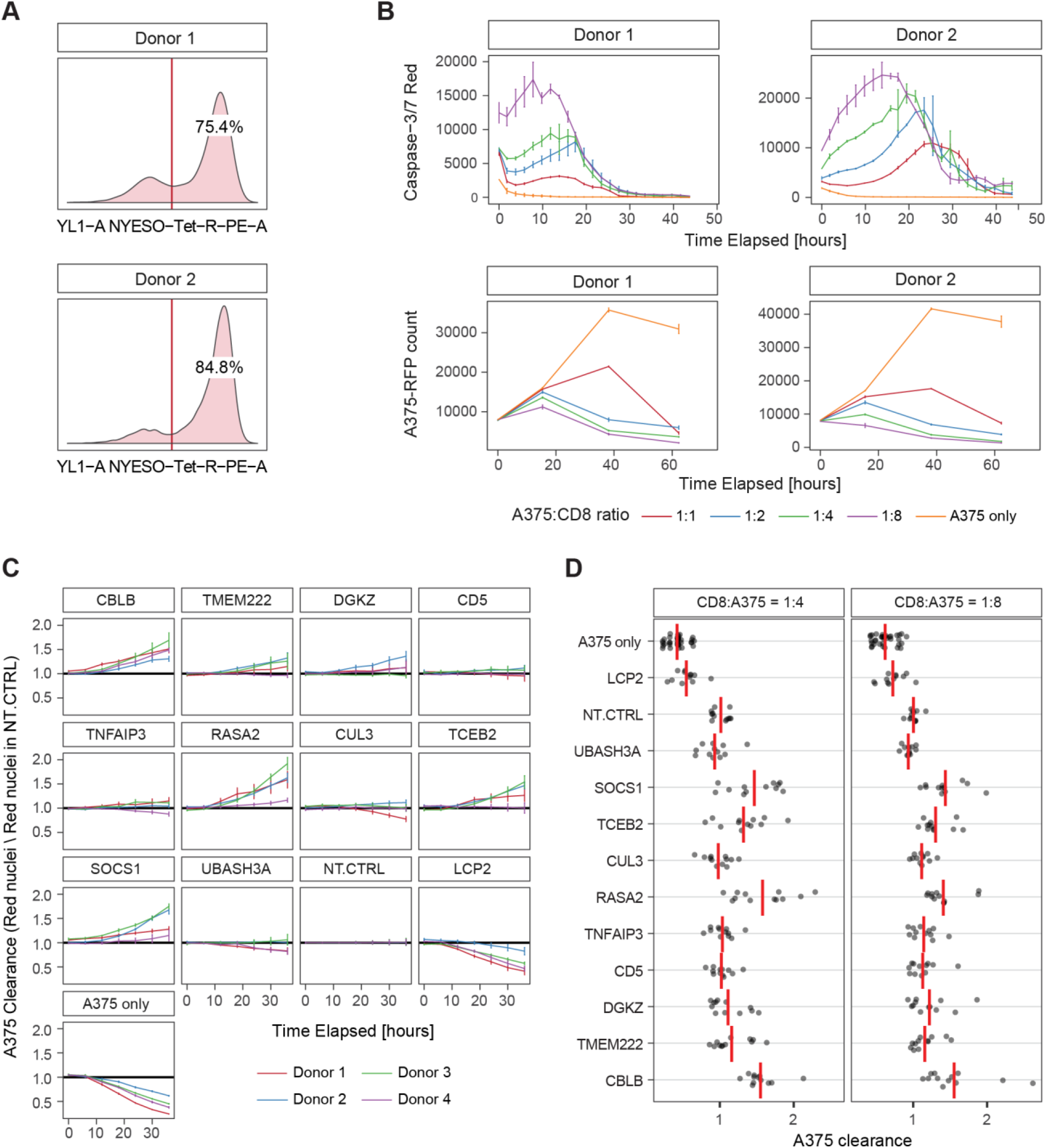
Cancer Cell Killing *In Vitro* by Engineered Human T Cells, Related to Figure 5. (A)T cells post transduction with the 1G4 TCR lentivirus showed high transduction rates based on staining with HLA-A2+ restricted NY-ESO-1 peptide dextramer. Cells were sorted to obtain a pure population of tetramer-positive, cancer specific CD8^+^ T cells. (B)Caspase levels measured on the IncuCyte demonstrate T cell-induced apoptosis of target NY-ESO expressing A375 melanoma cells in a titration of increasing T cell to tumor cell ratios (top panels). This cancer cell killing corresponded with the A375 RFP-tagged nuclear counts over time for the same T cell to cancer cell ratios (bottom panels). (C)A375 cell clearance by T cells with various target gene modifications. Y-axis shows A375 clearance – A375 nuclear count, normalized by the count in the NT.CTRL wells for each donor, time point, and gene target. Lines show the mean for each donor (n=4), across two guides and two technical replicates. Error bars are SEM. (D)Data in (B) quantified at 36 hours for two CD8 to A375 ratios. 1:4 is data summarized in Figure 5C. Dots represent individual wells in the arrays, two guides and two replicates, for all donors (n=4).

**Figure S6.**
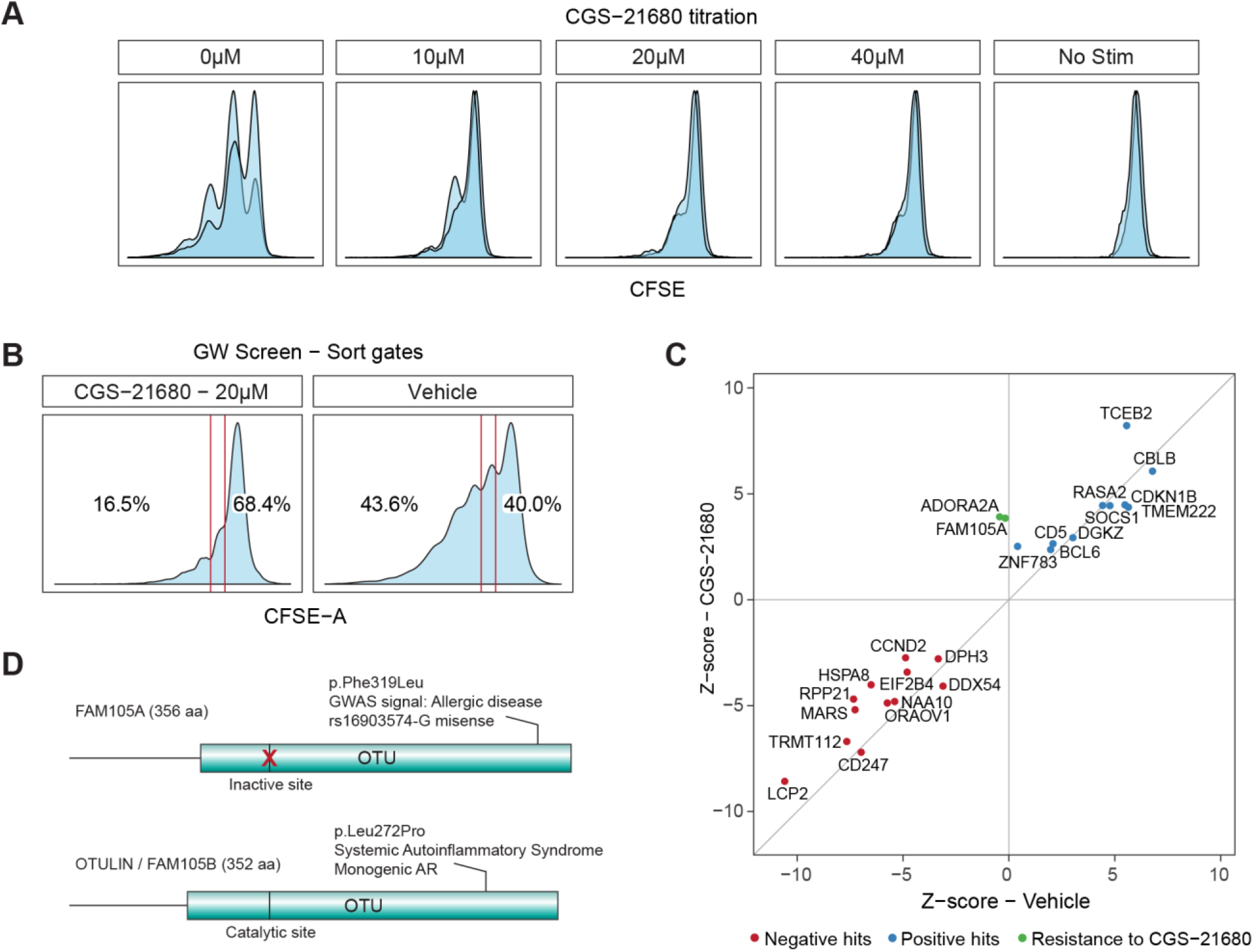
Screen for Resistance to an Immunosuppressive Signal in Primary T Cells Using SLICE, Related to Figure 6. (A)Dose titration of adenosine receptor agonist CGS-21680 for the optimal effects on T cell proliferation inhibition identified 20μM as sufficiently suppressive. Shown are representative CFSE profiles for varying concentrations of CGS-21680, as labeled at the top of each panel. Data is for CD8^+^ T cells from two human donors. (B)CFSE profile and sorting gates from the genome wide screen shown in Figure 6A. (C)Comparison of the overlapping top 25 hits for the two screen conditions (CGS-21680 and vehicle). We noted that while many hits from the vehicle screen also boosted proliferation in the presence of the immunosuppressive adenosine agonist (trending along diagonal), *ADORA2A* and *FAM105A* sgRNAs were enriched selectively in the setting of the adenosine agonist. *ADORA2A* and *FAM105A* were the only hits in the top 0.1% by LFC in the CGS-21680 treated condition and also in the top 0.1% by LFC difference between the CGS-21680 and the vehicle GW screen conditions. (D)Paralogs FAM105A and OTULIN (FAM105B) are 40% homologous and are encoded at neighboring sites on chromosome 5. They share an OTU deubiquitinase domain, which in FAM105A is predicted to be catalytically inactive. A non-synonymous single nucleotide polymorphism (SNP) in FAM105A has been linked to allergic disease by genome wide association studies (Ferreira et al., 2017). A rare mutation in OTULIN has been implicated in monogenic severe inflammatory disease (Damgaard et al., 2016).

